# Elucidation of the life cycle of a saprotrophic inoperculate discomycete that is associated with pinesap, using a polyphasic taxonomic approach^#^

**DOI:** 10.1101/2024.06.01.596929

**Authors:** Özge Demir, Yanpeng Chen, Christopher Lambert, Anja Schüffler, Frank Surup, Marc Stadler

**Affiliations:** Department Microbial Drugs, Helmholtz Centre for Infection Research (HZI), German Centre for Infection Research (DZIF), Partner Site Hannover-Braunschweig, Inhoffenstrasse 7, 38124 Braunschweig, Germany; Institute of Microbiology, Technische Universität Braunschweig, Spielmannstraße 7, 38106 Braunschweig, Germany; School of Life Science and Technology, University of Electronic Science and Technology of China, Chengdu, 611731, People’s Republic of China; Institut für Biotechnologie und Wirkstoff-Forschung gGmbH (IBWF), Hanns-Dieter-Hüsch-Weg 17, 55128 Mainz, Germany

**Keywords:** Monotropoid mycorrhiza, chemotaxonomy, secondary metabolites, *Leotiomycetes*, phylogeny

## Abstract

This study deals with the elucidation of the life cycle of an inoperculate discomycete that was eventually collected from conifer wood in South Germany and has become famous for the extraordinary diversity of its secondary metabolites when it was studied extensively during the 1990s. It had then been identified as *Lachnum papyraceum* (*Lachnaceae*, *Helotiales*) based on morphological traits, and extracts from its mycelial cultures were found to possess extraordinary nematicidal and antibiotic activities. Over 60 different secondary metabolites were finally identified from this fungus after extensive variation of culture media and scale up of production up to 100 litre scale. Among the main active principles were mycorrhizin A and chloromycorrhizin A, which had first been reported in 1987 from an unnamed “mycorrhizal” fungus of the hemiparasitic plant, *Monotropa hypopitys* (pinesap) that was isolated in Sweden. We noted that both the *Lachnum* strain and the original mycorrhizin producer were still available in the public domain, and decided to study them for comparison, using a multilocus phylogeny and also generated secondary metabolite profiles of both strains using analytical high performance liquid chromatography coupled to diode array and mass spectrometric detection (HPLC-DAD/MS). Surprisingly, the sequence data as well as the secondary metabolite profiles of both strains were highly similar, and it was also confirmed by phylogenetic methods that the strains are indeed nested within the genus *Lachnum* by comparison of their ITS, LSU and RBP2 sequences. The specimen called *L. papyraceum* in the old publications was tentatively re-identified by Hans-Otto Baral as *L*. cf. *subvirgineum,* but substantial further work on the taxonomy of the genus remains to be done, anyway. We conclude that some *Lachnum* species have a highly complex but all the more interesting life cycle, and the mycorrhizal symbiont partner may invade the host plant, where it may persist as an endophyte and finally turn saprotrophic on the wood of the senescent pine tree. The taxonomy of these fungi should also be further resolved in the future, using a polythetic concept that includes chemotaxonomic data and a multi-locus genealogy.

## Introduction

Fungi are nowadays very well-known to have great economical and ecological importance, as they play a pivotal role in both the global ecosystems and the global economy (Niego et al. 2023), but this has not always been the case. For instance, during the end of the last century, many academic and industrial research groups were working on the discovery of new biologically active secondary metabolites, based on screening of new isolates at random. During these activities, in which thousands of scientists were involved, many new metabolites with potentially beneficial activities have been discovered. However, producer strains were often not properly preserved, and not even characterized, using state of the art methods.

Only few pharmaceutical companies that were active in the past have maintained the necessary resources, especially as natural product screening eventually became out of fashion. Several large culture collections that had accumulated in the pharma industry have thus eventually become lost. An exception from this case, however, are so-called “patent strains” that found their way into the professional culture collections and other biodiversity repositories when the authors of patents needed to deposit the producer organisms as part of the patent application under the Budapest Treaty^1^ or other internationally valid agreements.

At the same time, the scientists in Academia who recovered producers of interesting compounds rarely had the foresight to deposit their producer strains in a public culture collection from where they could retrieved by others later on. Many of these studies were dominated by chemists, who did not care about fungal taxonomy or just did not possess the necessary resources to keep their fungal strains alive after the chemical work was done. Consequently, the only valuable information that remains until today are the chemical structures of these compounds and the corresponding analytical data.

Our current study is based on a lucky coincidence. We have not only been able to retrieve a strain that was previously deposited in the course of a patent application but even came across another one that was deposited by a Swedish academian who apparently recognized that he was dealing with something special and therefore sent his strain to the ATCC collection without any commercial interest (Trofast and Wickberg 1977).

During the literature review on our recent paper on a strain of the discomycete genus *Pezicula* that was antagonistic against the Ash dieback pathogen, *Hymenoscyphus fraxineus* (Demir et al 2023), we came across the highly potent antibiotic mycorrhizin A (**1**) that. This compound was identified as one of the main active principles of the antagonist, but at the same time, the fact that the fungus produced the toxic metabolite precluded its further development as biocontrol agent. In any case, we noted that among the previously reported sources for mycorrhizin A were two apparently unrelated fungal strains that were both still extant in public culture collections. Since we are presently working on the secondary metabolites of *Lachnum*, anyway (Phutthacharoen et al. 2023, 2024), we have ordered and studied these strains for secondary metabolite production as well as by state of the art molecular phylogenetic methods for the first time, to clarify whether they may actually be phylogenetically related to *Pezicula* (and/or or each other). The current paper is dedicated to answer this question.

## Material and methods

### General experimental procedures

#### Fungal material

The strains studied were *Lachnum papyraceum* A48-88 (Stadler et al. 1993, deposited as DSM 10201 with the DSMZ culture collection, Braunschweig, Germany) and ATCC 36554 (deposited as an unnamed “mycorrhizal fungus D37” with the American Type Culture Collection, Manassas, VA, USA). The latter culture had been obtained from roots of *Monotropa hypopitys i*n Sweden (Trofast and Wickberg 1987). The strain DSM 10201 was originally isolated from a specimen collected by by Heidrun Anke in Hinterstein, Germany, in 1988, and identified by Wolf-Rüdiger Arendholz. A voucher specimen was deposited at the university of Kaiserslautern, and is now housed at the IBWF in Mainz. According to an examination by Hans-Otto Baral, it is actually more closely related to *L. subvirgineum*, according to the current taxonomic concept. The exact classification of the fungus in comparison with type material will be reported separately as it is beyond the scope of the current study, and *L. subvirgineum* is not even validly described. We just refer to the specimen A 48-88 here as *L.* cf. *subvirgineum*.

#### Fermentation and extraction

For comparison of secondary metabolite production, strains ATCC 36554 and *Lachnum* cf. *subvirgineum* strain DSM 10201 were fermented as previously described by Stadler et al. (1993) in MGP medium.

The cultures were incubated on a rotary shaker at 23 °C and 140 min^−1^. The glucose consumption was monitored by constantly checking the amount of free glucose using Medi-test Glucose (Macherey-Nagel, Düren, Germany). Glucose was depleted after six days of cultivation for strain ATCC 36554 and after twenty days for DSM 10201. The fermentation was terminated 4 days after glucose depletion. Subsequently, the mycelium and supernatant were separated via vacuum filtration. The mycelia were then extracted twice with acetone in an ultrasonic bath at 40 °C for 30 min. The resulting acetone extracts were dried in vacuo at 40°C. The remaining aqueous residues were diluted with the same amounts of ethyl acetate and extracted one time. The supernatants were extracted twice with the same amount of EtOAc. The solvents were then dried in vacuo at 40 °C.

#### Analytical HPLC

The extracts of both strains were dissolved with methanol to yield a concentration of 4.5 mg/mL.

HPLC-DAD/MS analyses were performed on the amaZon speed ETD ion trap mass spectrometer (Bruker Daltonics, Bremen, Germany) in both positive and negative ionization modes simultaneously and HR-ESI-MS (high-resolution electrospray ionization mass spectrometry) analyses were measured on the MaXis ESI-TOF (time of flight) mass spectrometer (Bruker Daltonics) coupled to an Agilent 1260 series HPLC-UV system (Agilent Technologies, Santa Clara, CA, USA). Following the injection of two microliters of the samples, the separation was carried out with an ACQUITY-UPLC BEH C18 column (50 × 2.1 mm; particle size: 1.7 μm) by Waters. HPLC grade water and HPLC grade acetonitrile supplemented by 0.1% formic acid were used as mobile phase. The elution gradient started at 5% of acetonitrile, continued with an increase to 100% in 20 min and retained at 100% for 5 min. The flow rate was 0.6 mL/min, and the UV/Vis detections were recorded at 190–600 nm and 210 nm. The chromatograms were analyzed using the software Data Analysis 6.1 (Bruker) and the spectra were compared to an internal database that contained standards of known secondary metabolites from prior studies.

#### Molecular phylogeny

The DNA sequence data of both strains (ATCC 36554 and DSM 10201) were generated as described in details by Wendt et al. (2018) using exactly the same equipment. Based on previous studies (Han et al. 2014, Hongsanan et al. 2015; for details see Table 5), in total 16 ITS, 15 LSU and 15 *RPB2* sequences of *Lachnum* species and closely related taxa were obtained from GenBank. *Arachnopeziza aurelia* (TNS-F11211) and *A. aurata* (TNS-F11212) were selected as the outgroup taxa. The multiple sequence alignments were performed using MAFFT version 7.310 (Katoh et al 2002) followed by additional trimming of the alignment results using trimAl version 1.4 (Capella-Gutiérrez et al. 2009) with the “-gapthreshold 0.5” option, which allows only 50% of taxa with a gap in each site. Maximum likelihood (ML) phylogenetic trees were obtained using RAxML-NG version 1.2.1 (Kozlov et al 2019), and the resulting topology was assessed through 1000 bootstrap replicates. GTR+I+G was chosen as the best-fit model for LSU, *RPB2* and ITS. The tree was visualized using ggtree (Yu 2020) and edited in Adobe Illustrator version 20.0.0. The reference sequences and their origin are summarized in Table 1.

**Table 1.** Taxa information used in the present study and their corresponding GenBank accession numbers.

| Species | Specimen<br>/Isolate no. | GenBank Accession number |  |  | References |
| --- | --- | --- | --- | --- | --- |
|  |  | ITS | LSU | *RPB2* |  |
| *Lachnum rachidicola* | TNS:F-81229<br>= NBRC<br>114473 | LC669467 | LC533136 | LC533189 | Tochihara and<br>Hosoya 2022 |
| *Lachnum virgineum* | TNS:F-16583 | AB481268 | AB481306 | AB481343 | Hosoya et al<br>2010 |
| *Lachnum soppittii* | FC-2160 | AB481266 | AB481308 | AB481344 | Hosoya et al<br>2010 |
| *Lachnum controversum* | MFLU 18-1820 | MK584937 | MK591964 | MK368613 | Ekanayaka et<br>al 2019 |
| “*Lachnum*”<br>*palmae* | TNS: F-24600 =<br>NBRC<br>114443 | LC669444 | LC533145 | LC533224 | Tochihara and<br>Hosoya 2022 |
| “*Lachnum*<br>*abnorme*”<br>(currently valid<br>name:<br>*Erioscyphella*<br>*abnormis*) | KUS-F52080 | JN033395 | JN086698 | JN086849 | Han et al<br>2014<br>Han et al<br>2021 |
| *Trichopezizella*<br>sp. | KUS-F52478 | JN033417 | JN086720 |  | Han et al<br>2014a |
| *Trichopeziza*<br>*sulphurea* | KUS-F52218 | JN033398 | JN086701 | JN086852 | Han et al<br>2014a |
| “*Lachnum*<br>*fuscescens*”<br>(currently valid<br>name:<br>*Brunnipila*<br>*fuscescens*) | FC-2200 | AB481255 | AB481311 | AB481348 | Hosoya et al<br>2010 |
| *Brunnipila*<br>*fuscescens* | KUS-F52031 | JN033392 | JN086695 | JN086846 | Han et al<br>2014a |
| *Proliferodiscus*<br>*inspersus* var.<br>*magniascus* | KUS-F52660 | JN033427 | JN086730 | JN086871 | Han et al<br>2014a,b |
| *Proliferodiscus*<br>*inspersus* var.<br>*magniascus* | TNS: F-17436 | JN033452 | JN086752 | JN086898 | Han et al<br>2014a,b |
| *Arachnopeziza*<br>*aurelia* | TNS: F-11211 | JN033435 | AB546937 | JN086880 | Han et al<br>2014a,b |
| *Arachnopeziza*<br>*aurata* | TNS: F-11212 | JN033436 | AB546936 | JN086881 | Han et al<br>2014a,b |
| *Lachnum*<br>*papyraceum* | DSM 10201 | PP702451 | PP698021 | PP711689 |  |
| *Lachnum* sp. | ATCC 36554 | PP702452 | PP698022 | PP711690 |  |

**Table 2.**
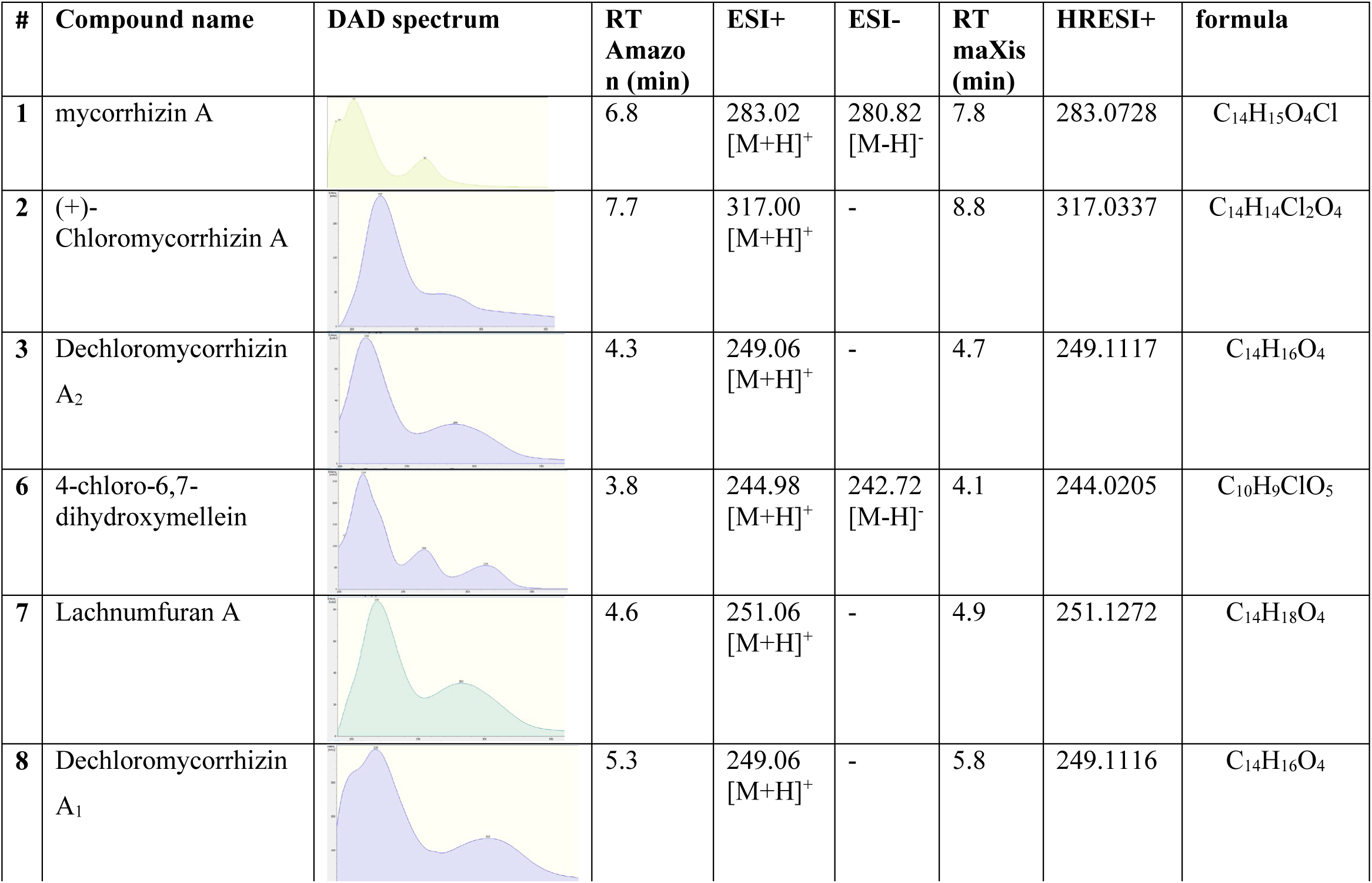
List of identified compounds of crude extracts from the supernatant of *Lachnum* sp. (ATCC 36554). DAD spectra, Rt (retention time) and masses were indicated.

| # | Compound name | DAD spectrum | RT<br>Amazo<br>n (min) | ESI+ | ESI- | RT<br>maXis<br>(min) | HRESI+ | formula |
| --- | --- | --- | --- | --- | --- | --- | --- | --- |
| 1 | mycorrhizin A |  | 6.8 | 283.02<br>[M+H]<sup>+</sup> | 280.82<br>[M-H]<sup>-</sup> | 7.8 | 283.0728 | C<sub>14</sub>H<sub>15</sub>O<sub>4</sub>Cl |
| 2 | (+)-<br>Chloromycorrhizin A |  | 7.7 | 317.00<br>[M+H]<sup>+</sup> | - | 8.8 | 317.0337 | C<sub>14</sub>H<sub>14</sub>Cl<sub>2</sub>O<sub>4</sub> |
| 3 | Dechloromycorrhizin<br>A<sub>2</sub> |  | 4.3 | 249.06<br>[M+H]<sup>+</sup> | - | 4.7 | 249.1117 | C<sub>14</sub>H<sub>16</sub>O<sub>4</sub> |
| 6 | 4-chloro-6,7-dihydroxymellein |  | 3.8 | 244.98<br>[M+H]<sup>+</sup> | 242.72<br>[M-H]<sup>-</sup> | 4.1 | 244.0205 | C<sub>10</sub>H<sub>9</sub>ClO<sub>5</sub> |
| 7 | Lachnumfuran A |  | 4.6 | 251.06<br>[M+H]<sup>+</sup> | - | 4.9 | 251.1272 | C<sub>14</sub>H<sub>18</sub>O<sub>4</sub> |
| 8 | Dechloromycorrhizin<br>A<sub>1</sub> |  | 5.3 | 249.06<br>[M+H]<sup>+</sup> | - | 5.8 | 249.1116 | C<sub>14</sub>H<sub>16</sub>O<sub>4</sub> |

**Table 3.**
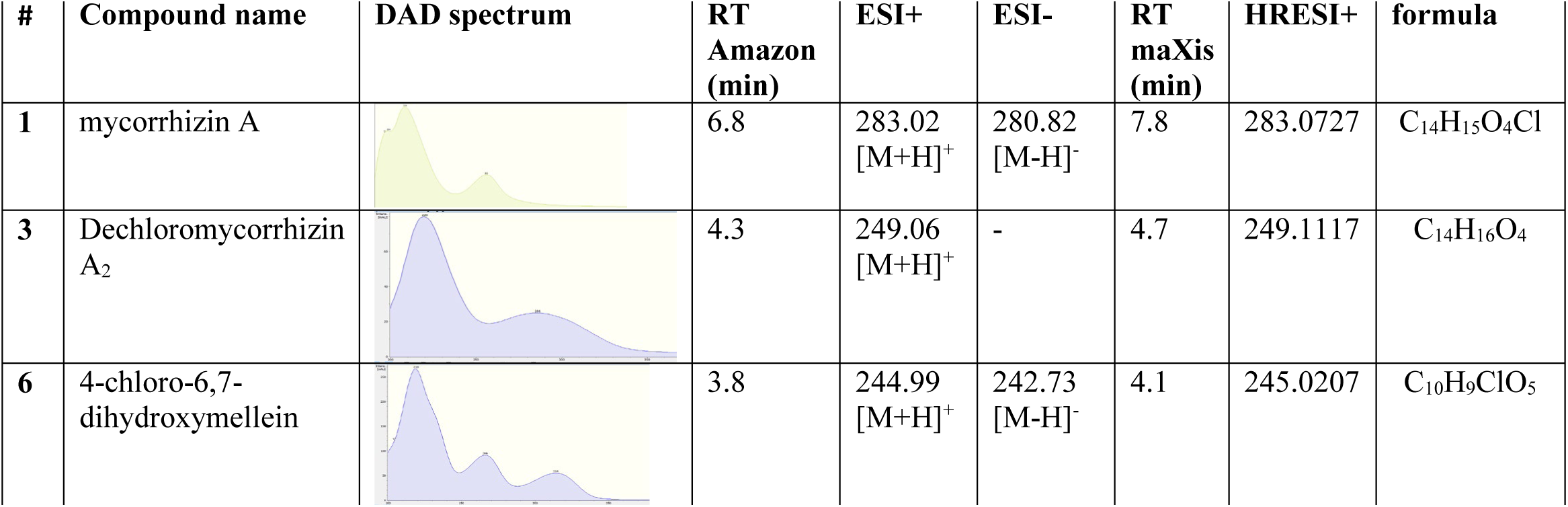
List of identified compounds of crude extracts from the mycelium of *Lachnum* sp. (ATCC 36554). DAD spectra, Rt (retention times) in two different HPLC systems and masses were indicated.

| # | Compound name | DAD spectrum | RT<br>Amazon<br>(min) | ESI+ | ESI- | RT<br>maXis<br>(min) | HRESI+ | formula |
| --- | --- | --- | --- | --- | --- | --- | --- | --- |
| 1 | mycorrhizin A |  | 6.8 | 283.02<br>[M+H]<sup>+</sup> | 280.82<br>[M-H]<sup>-</sup> | 7.8 | 283.0727 | C<sub>14</sub>H<sub>15</sub>O<sub>4</sub>Cl |
| 3 | Dechloromycorrhizin<br>A<sub>2</sub> |  | 4.3 | 249.06<br>[M+H]<sup>+</sup> | - | 4.7 | 249.1117 | C<sub>14</sub>H<sub>16</sub>O<sub>4</sub> |
| 6 | 4-chloro-6,7-dihydroxymellein |  | 3.8 | 244.99<br>[M+H]<sup>+</sup> | 242.73<br>[M-H]<sup>-</sup> | 4.1 | 245.0207 | C<sub>10</sub>H<sub>9</sub>ClO<sub>5</sub> |

**Table 4.**
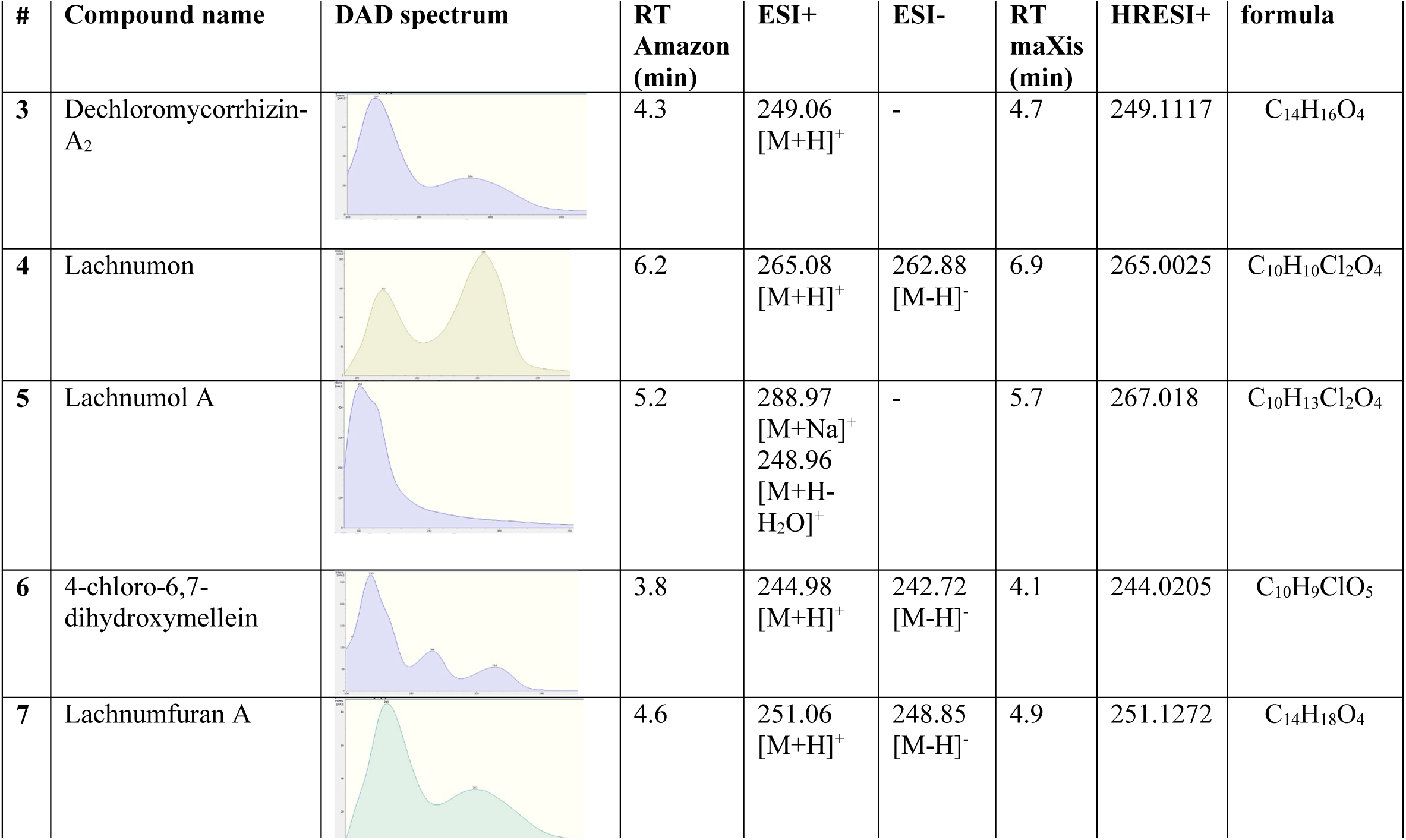
List of identified compounds of crude extracts from the supernatant of *Lachnum papyraceum* (DSM 1020). DAD spectra, Rt (retention time) and masses were indicated.

| # | Compound name | DAD spectrum | RT<br>Amazon<br>(min) | ESI+ | ESI- | RT<br>maXis<br>(min) | HRESI+ | formula |
| --- | --- | --- | --- | --- | --- | --- | --- | --- |
| 3 | Dechloromycorrhizin-A<sub>2</sub> |  | 4.3 | 249.06<br>[M+H]<sup>+</sup> | - | 4.7 | 249.1117 | C<sub>14</sub>H<sub>16</sub>O<sub>4</sub> |
| 4 | Lachnumon |  | 6.2 | 265.08<br>[M+H]<sup>+</sup> | 262.88<br>[M-H]<sup>-</sup> | 6.9 | 265.0025 | C<sub>10</sub>H<sub>10</sub>Cl<sub>2</sub>O<sub>4</sub> |
| 5 | Lachnumol A |  | 5.2 | 288.97<br>[M+Na]<sup>+</sup><br>248.96<br>[M+H-H<sub>2</sub>O]<sup>+</sup> | - | 5.7 | 267.018 | C<sub>10</sub>H<sub>13</sub>Cl<sub>2</sub>O<sub>4</sub> |
| 6 | 4-chloro-6,7-dihydroxymellein |  | 3.8 | 244.98<br>[M+H]<sup>+</sup> | 242.72<br>[M-H]<sup>-</sup> | 4.1 | 244.0205 | C<sub>10</sub>H<sub>9</sub>ClO<sub>5</sub> |
| 7 | Lachnumfuran A |  | 4.6 | 251.06<br>[M+H]<sup>+</sup> | 248.85<br>[M-H]<sup>-</sup> | 4.9 | 251.1272 | C<sub>14</sub>H<sub>18</sub>O<sub>4</sub> |

**Table 5.**
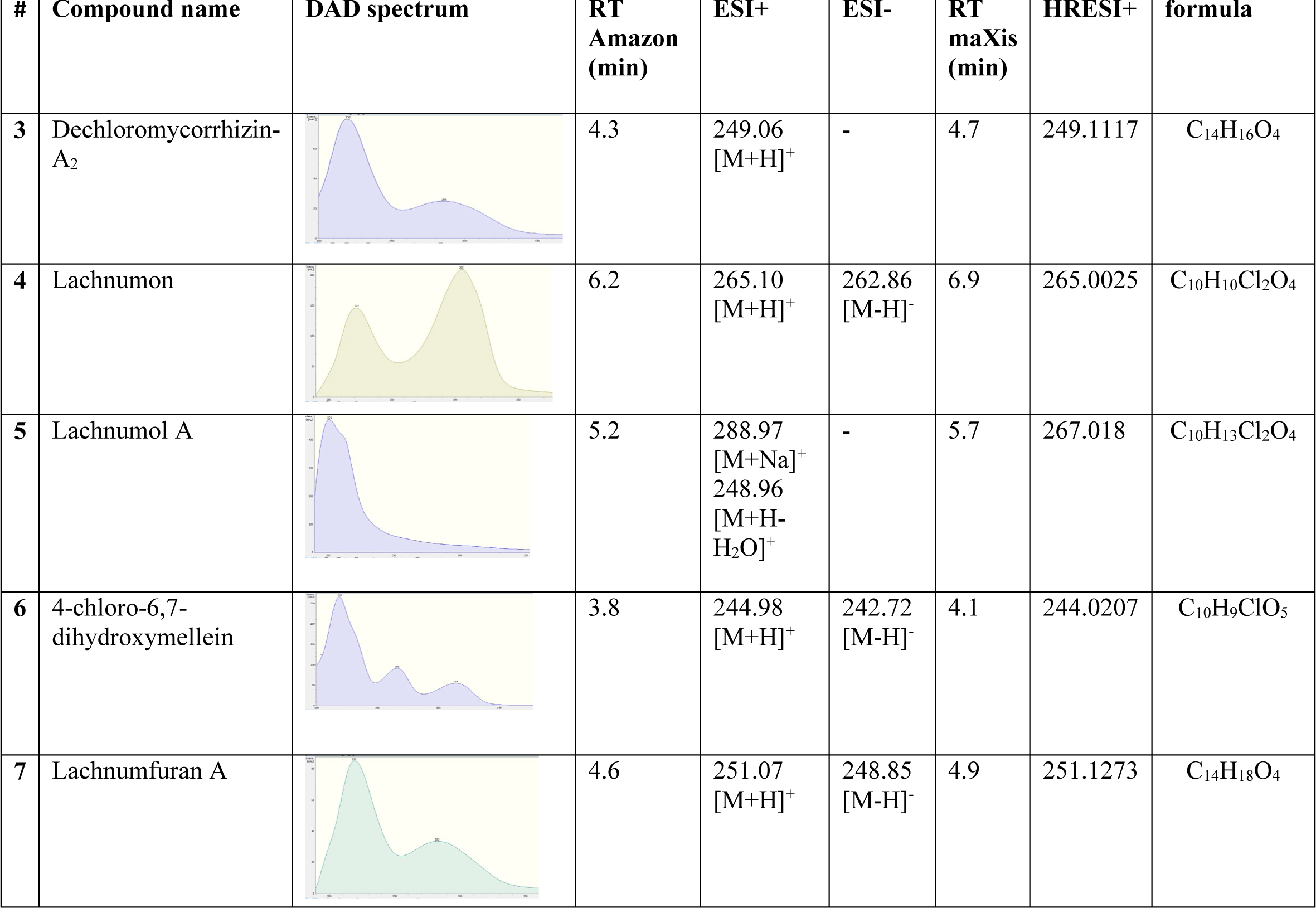
List of identified compounds of crude extracts from the mycelium of *Lachnum papyraceum* (DSM 1020). DAD spectra, Rt (retention time) and masses were indicated.

| # | Compound name | DAD spectrum | RT Amazon (min) | ESI+ | ESI- | RT maXis (min) | HRESI+ | formula |
| --- | --- | --- | --- | --- | --- | --- | --- | --- |
| 3 | Dechloromycorrhizin-A<sub>2</sub> |  | 4.3 | 249.06 [M+H]<sup>+</sup> | - | 4.7 | 249.1117 | C<sub>14</sub>H<sub>16</sub>O<sub>4</sub> |
| 4 | Lachnumon |  | 6.2 | 265.10 [M+H]<sup>+</sup> | 262.86 [M-H]<sup>-</sup> | 6.9 | 265.0025 | C<sub>10</sub>H<sub>10</sub>Cl<sub>2</sub>O<sub>4</sub> |
| 5 | Lachnumol A |  | 5.2 | 288.97 [M+Na]<sup>+</sup><br>248.96 [M+H-H<sub>2</sub>O]<sup>+</sup> | - | 5.7 | 267.018 | C<sub>10</sub>H<sub>13</sub>Cl<sub>2</sub>O<sub>4</sub> |
| 6 | 4-chloro-6,7-dihydroxymellein |  | 3.8 | 244.98 [M+H]<sup>+</sup> | 242.72 [M-H]<sup>-</sup> | 4.1 | 244.0207 | C<sub>10</sub>H<sub>9</sub>ClO<sub>5</sub> |
| 7 | Lachnumfuran A |  | 4.6 | 251.07 [M+H]<sup>+</sup> | 248.85 [M-H]<sup>-</sup> | 4.9 | 251.1273 | C<sub>14</sub>H<sub>18</sub>O<sub>4</sub> |

**Table 6.**
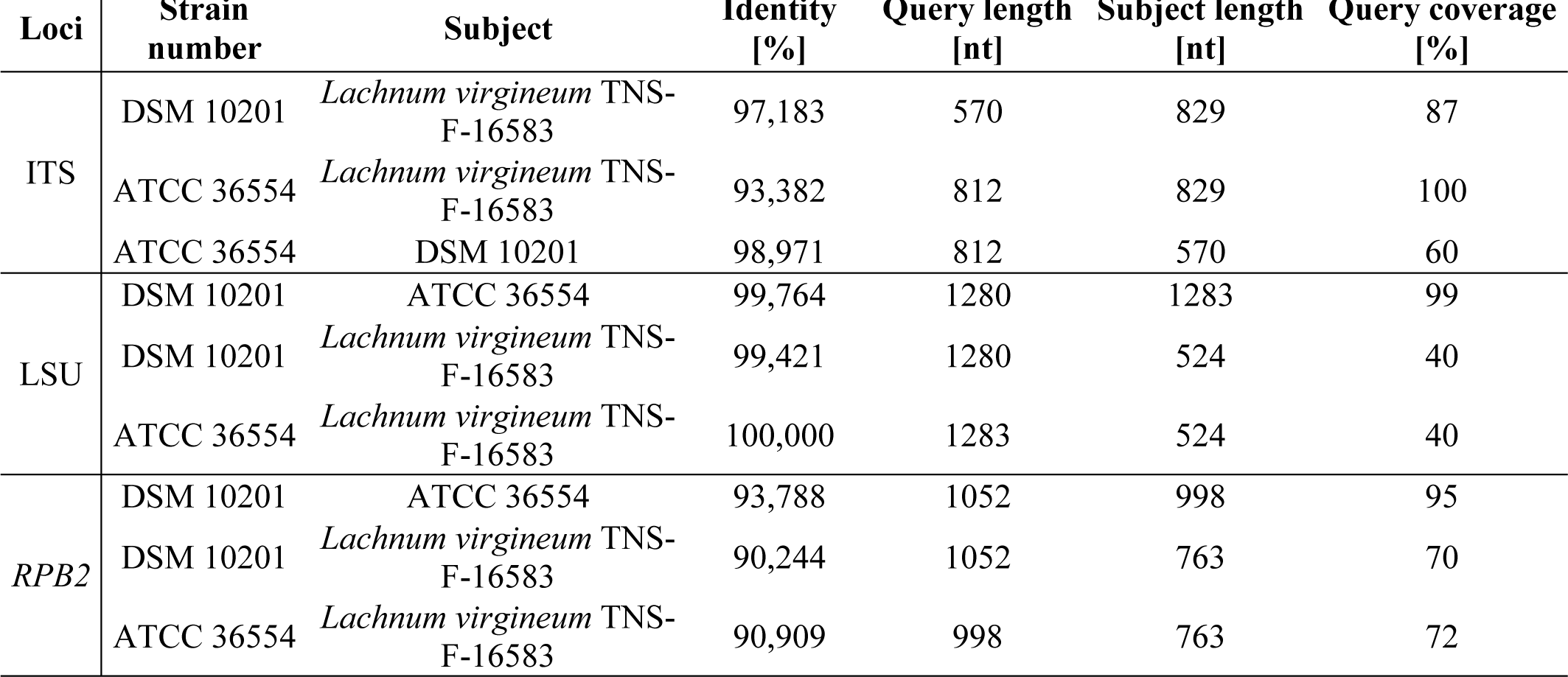
Pairwise sequence identities for three loci from BLAST analysis.

## Results and Discussion

The two strains under study had been previously reported to produce a rare family of bioactive metabolites that has apparently so far only been encountered in cultures of species belonging to the *Helotiales* (Schulz et al.1995; McMullen et al. 2017; Demir et al. 2023). While only the mycorrhizins were reported from the ATCC strain, which also produced a different derivative named chloromycorrhizinol A (Trofast 1978) strain DSM 10201 has been studied rather extensively in the past. Aside from culture media variations, even a scale-up to 100 liter scale was carried out. Some of the results have been summarized by Anke et al. (1995) but some follow-up studies yet yielded further metabolites and some of those were even patented by Hannske et al. (1999). We have therefore used the optimal medium reported for secondary metabolite production by Stadler et al. (1993) for both strains and found that they had very similar growth characteristics and gross morphology (see Fig. 1 for the colonies on agar plates).

**Figure 1.**
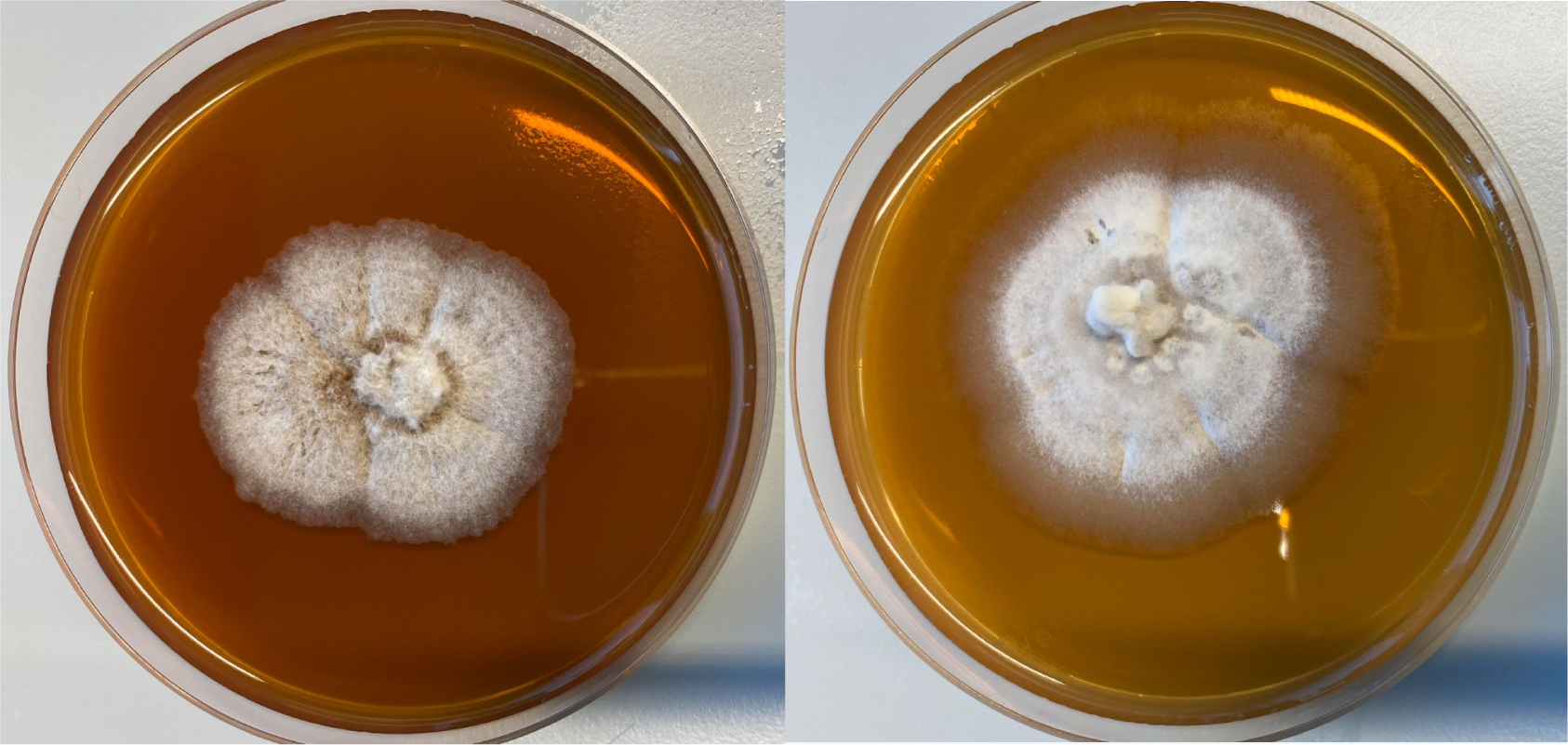
Three weeks growth of strain ATCC 36554 (left) and *Lachnum papyraceum* (DSM 1020) (right) on YM medium at 23 °C.

**Figure 2.**
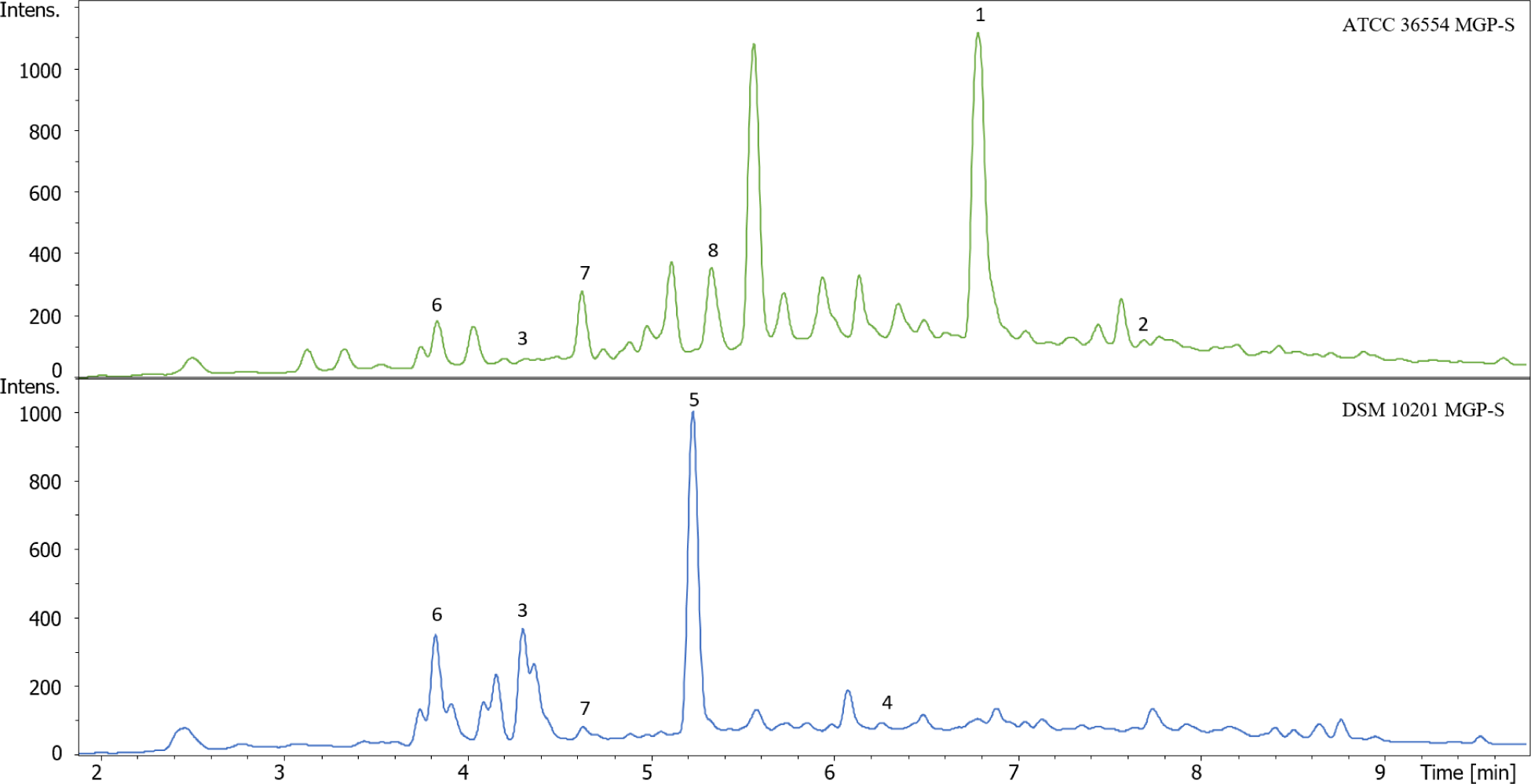
HPLC-UV chromatograms (210 nm) of the crude extracts from the supernatant of strains ATCC 36554 and DSM 10201 in MGP medium.

Even the secondary metabolite profiles of both cultures were highly similar, and aside from the mycorrhizins (**1** and **2**) several other compounds that had previously been reported from *L. papyraceum* were also prevalent in the other strain (see Fig. 3 and Table 1).

**Figure 3:**
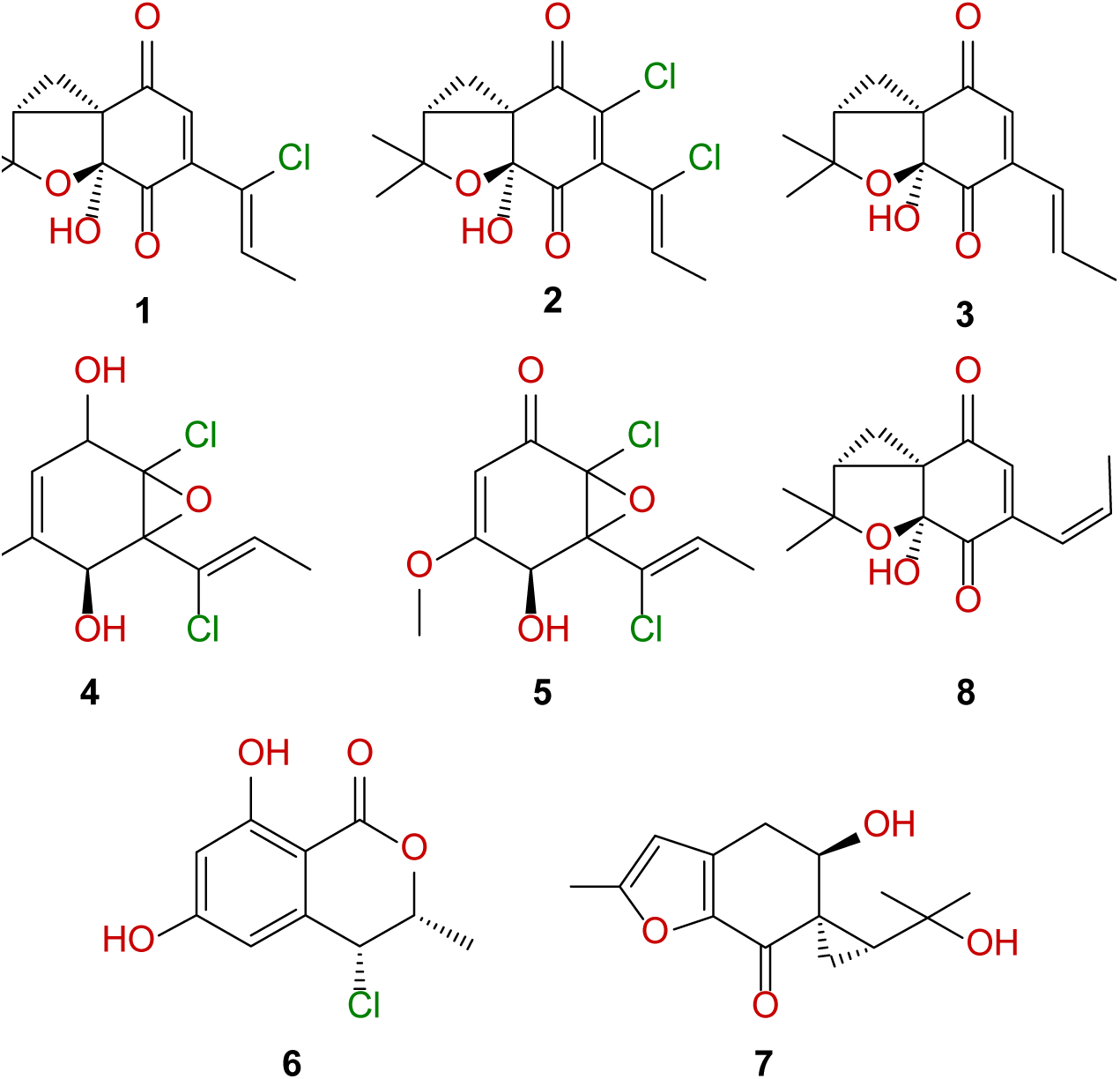
Chemical structures of the detected compounds. **1**: Mycorrhizin A; **2**: Chloromycorrhizin A; **3**: Dechloromycorrhizin A1; **4**: Lachnumol A; **5**: Lachnumon; **6**: 4-chloro-6,7-dihydroxymellein; **7:** Lachnumfuran A; **8**: Dechloromycorrizin A2.

Strain ATCC 36554 produced mycorrhizin A (**1**) as major compound. Chloromycorrhizin A (**2**), dechloromycorrhizin-A (**3**) and 4-chloro-6,7-dihydroxymellein (**6**) were detected as minor metabolites. On the other hand, strain DSM 10201 contained lachnumol A (**5**) as the major metabolite along with lachnumon (**4**) and 4-chloro-6,7-dihydroxymellein (**6**) as minor compounds. Both strains produced dechloromycorrhizin-A (**3**) and 4-chloro-6,7-dihydroxymellein (**6**). Lachnumfuran A (**7**) was also tentatively detected in both strains. These chemotaxonomic data strongly point toward a very close relationship of both strains.

The phylogram (Fig. 3) also clearly shows that the strains belong to the genus *Lachnum*. The combined dataset included three loci, namely ITS, LSU, and *RPB2*, from 14 isolates of *Lachnum* s.l. with *Arachnopeziza aurelia* (TNS-F11211) and *A. aurata* (TNS-F11212) as the outgroup taxa.

Blast results indicated that the best hit for DSM 10201 was “uncultured Helotiales isolate B105_4” with sequence identity of 99.1% (0 gap) using ITS sequence. Similarly, the best hit for ATCC 36554 was “*Lachnum* sp. strain UNIJAG.PL.686” with sequence identity of 99.1% (0 gap) using the ITS sequence. For LSU, the best hit was “*Lachnum* cf. *pygmaeum* isolate NB-175-2” with sequence identity 99.5% (0 gap), and for ATCC 36554, the best hit was also “*Lachnum* cf. *pygmaeum* isolate NB-175-2” with sequence identity 99.8% (0 gap). In the case of *RPB2*, the best hit for both DSM 10201 and ATCC 36554 was “*Lachnum virgineum* isolate AFTOL-ID 49” with a sequence identities of 90.8% (4 gaps) and 90.4% (4 gaps), respectively.

RAxML analysis yielded a best-scoring tree (Fig. 4) with a final ML optimization likelihood value of -10742.410675. The parameters were as follows: A= 0.256413, C= 0.218439, G= 0.270162, T= 0.254986; AC= 1.558842, AG= 3.459272, AT= 1.428661, CG= 0.997875, CT= 8.125349, GT= 1.000000; Gamma distribution shape parameter α = 0.194413; Tree-Length= 1.388600.

**Figure 4.**
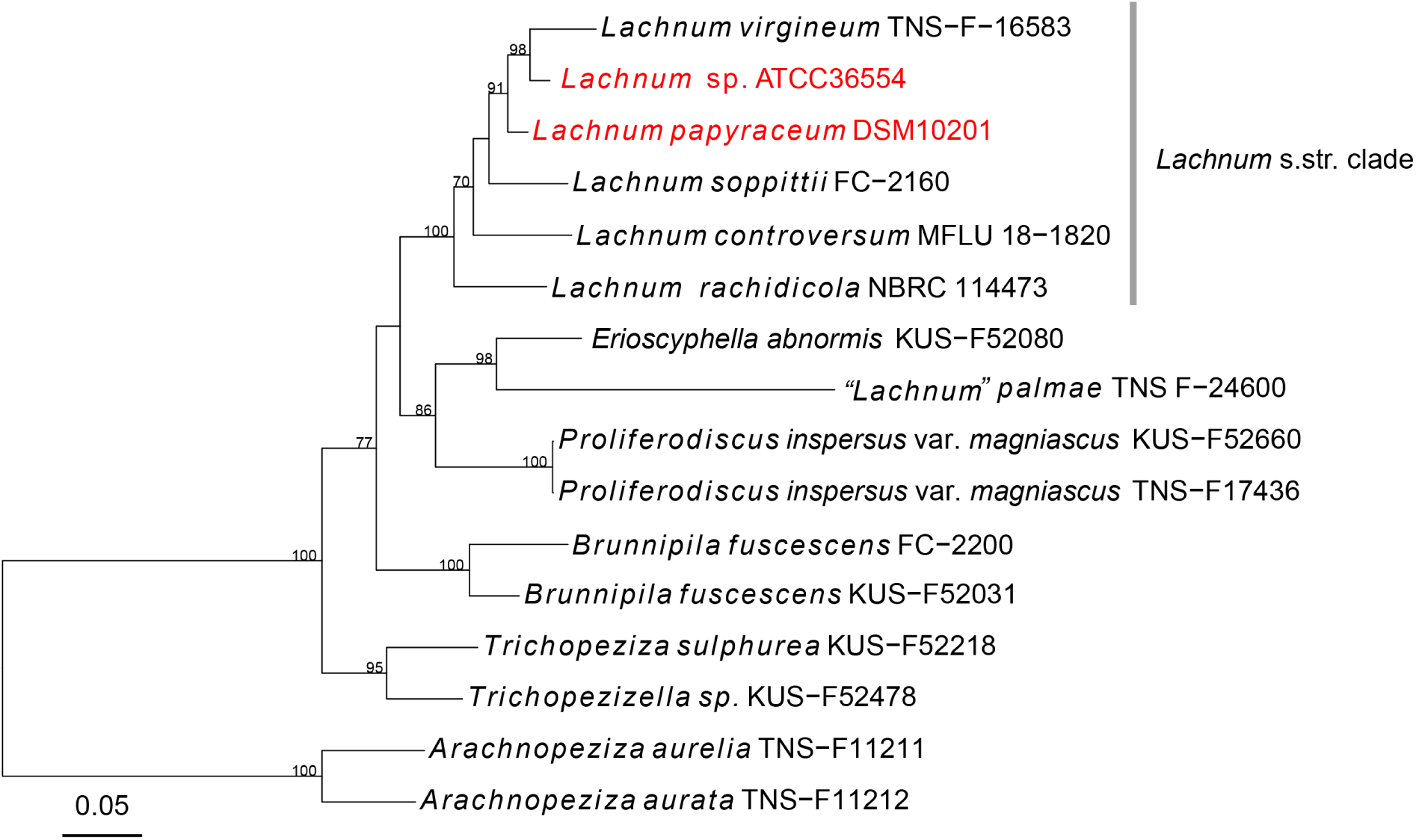
RAxML phylogram obtained from the combined ITS, LSU and *RPB2* sequences of the strains of *Lachnum* and its allied fungi. The strains ATCC 36554 and DSM 10201, which are discussed in this study, are highlighted in red. RAxML-NG bootstrap support (BS) values above 70% is shown at the nodes.

The best matches to the sequences of our two isolates were assigned to different epithets or not identified to the species level. The corresponding specimens do not represent type material and the corresponding cultures, if any were obtained, do not seem to be extant in public collections. Even if the specimens were still extant it would be impossible to generate data to assess the chemotaxonomic relationships because these discomycetes must be cultured from freshly collected material. The best match of our *RPB2* sequences was from “*Lachnum virgineum* isolate AFTOL-ID 49”, a specimen derived from the **“**Assembling The Fungal Tree Of Life” project that should be correctly identified. Nevertheless the data at hand only allow for assignment of the two mycorrhizin producing strains to the genus level. To date, there are a total of 532 records of names in *Lachnum* fide Index Fungorum (http://www.indexfungorum.org, accessed in March 2023) but sequence data are not available for most of them, hampering the species identification using molecular data. The multilocus phylogeny confirmed that our isolates belong to *Lachnum* and are very close to *Lachnum virgineum*. Our two isolates DSM 10201 and ATCC 36554 showed only >97 % and >93 % sequence identities in ITS with *Lachnum virgineum* TNS-F-16583, and >90 %, and >90 % in *RPB2* respectively, strongly suggesting that they are different species. Additionally, our two isolates can be distinguished by the rather low sequence identity (>93 %) in *RPB2*. It often pays off to sequence additional DNA loci if rDNA cannot give conclusive results.

In summary, all the data that we have gathered point toward the “mycorrhizal” fungus that was used in the study of Trofast and Wickberg (1977) being closely related, if not identical to the *Lachnum papyraceum* strain that was studied by the Anke group in the 1980s and 1990s.

Taxonomic concepts in the early 1990s were exclusively based on teleomorphic morphology, and no detailed description of the microscopic characteristics were reported in the earlier papers. In addition, even the morphology of the inoperculate discomycetes has since then undergone a rather revolutionary development, since Hans-Otto Baral and other workers started to use the so-called “Vital Taxonomy” to characterize these fungi over 40 years ago (cf. Baral 1992 and some examples for treatments of *Helotiales* by Baral et al. 2012; 2013 and Quijada et al. 2014). While these studies have certainly provided a lot of new evidence about the diversity of this largely neglected, yet ecologically interesting and highly important group of *Ascomycota*, there are some major drawbacks.

- Despite the descriptions often included morphological details of the morphological features and in some cases, even anamorph traits, fungi were often not kept “vital” -in many cases no cultures were obtained and preserved in public domain repositories.
- Ancient type specimens are often lost, or even if they are still extant, they are highly depauperate, and they are dead in any case. It is often not possible to assess the salient characters that serve for discrimination of species based on living material by direct comparison with the old types.
- The few available studies that attempted to link molecular data with vital taxonomy are unfortunately solely based on rDNA, loci like ITS and LSU. These loci are known to be very valuable for barcoding (as even the current study shows), and were generally employed for taxonomy in the early stages of molecular fungal taxonomy. However, they have later proven to be of little value in many of the taxonomic groups of *Ascomycota* (e.g., the entire classes of *Dothideomycetes*, *Eurotiomycetes*, and *Sordariomycetes*) where sufficient data eventually became available.
- Another problem that persists is the fact that relatively few *Leotiomycetes* have been cultured and studied for anamorphic characters, even though (as even this study shows) it is often possible to get their ascospores to grow in artificial culture media.

Interestingly, McMullin et al. (2017) reported mycorrhizins from a *Picea* inhabiting fungus that they tentatively classified as *Lachnum* cf. *pygmaeum*. Only an ITS sequence (GenBank acc no KY200575) was apparently used to aid classification, but no morphological data were reported. That species is also known to form apothecia on conifers, but differs from *L. papyraceum* in morphological traits of the teleomorph, and the anamorphic features are unknown for both species as far as we are aware^2^. The identification seems to have been based on an ITS sequence with GenBank acc. no OM456203, which was deposited by Hans-Otto Baral. It is an ascospore isolate from apothecia growing on an *Ulex* branch. The identity of this sample as *L. pygmaeum* was based on apothecial morphology (H-O. Baral, pers. comm.).

There are many other sequences in GenBank under the name *L. pygmaeum,* including e.g. MT276007 that was generated by Ramsfield et al. (2020) in the course of a molecular ecology study from a sample of the rhizosphere of aspen, and various others originating from China. However, there are currently no entries in GenBank (accessed on April 24, 2025) under the name *L. papyraceum*. A look at the BLAST search of our DSM strain of *L. papyraceum* reveals that most of the similar sequences that are already available were derived from molecular ecology studies and often not even assigned to a genus, or they were generated by amateur mycologists who depicted the images of the apothecia on the iNaturalist^3^ website (see Table S1 for the origin of the ten best matches). Not a single one of these sequences goes back to a fungal culture that could be studied for production of the mycorrhizins and other metabolites that are treated in the current study. On the other hand, it was interesting to observe that some of the closest matches were even derived from soil in grassland without any mention of conifers or *Monotropa* species. This would suggest that fungi phylogenetically related to *L. papyraceum* can also colonize totally different habitats and their life cycle may thus be even more complex than shown by the current study.

These observations are only included here to illustrate how much work needs to be done before the taxonomic status of our fungi, the distribution of mycorrhizins and other unique secondary metabolites that are known from *Lachnum* can be clarified. It should first be attempted to do some actual type studies of the old herbarium specimens, and see if they still bear the important diagnostic features that were recognized only a century later.

In any case, solving all these riddles and in particular the typification matters go beyond the scope of our current study, but we hope that our contribution can encourage other researchers to conduct follow-up work. In particular, more specimens of these tiny cup fungi should be cultured, so they can be studied for secondary metabolite production, anamorphic traits, molecular phylogeny based on multiple DNA loci, and ultimately even phylogenomics.

In this context, it is worthwhile to mention that Fehrer et al. (2019) have already reported a similar case where *Leotiomycetes* that are associated with *Ericaceae* are frequently encountered as endophytes. This study had been very helpful to solve many questions about the ecology and taxonomy of the fungi involved. The work presented here, however, can only be a first step towards a larger study involving more extensive sampling from both, the roots of *Monotropa* species and gaining cultures from the apothecia of the conifer-inhabiting cup fungi.

## Conclusions

The current study has revealed exciting details about the complex ecology of inoperculate discomycetes, which should give reason for various follow-up studies. After all, and as exemplified by the *Lachnum* cf. *subvirgineum* strain, these fungi have developed an extraordinarily rich secondary metabolism and this may well be due to their complex lifestyle, where they are exposed to various interorganismic interactions. It should be interesting to try to culture more of these fungi (in particular those that have been detected using ITS barcoding in various habitats but could not even be assigned to a genus for lack of reference data). While ITS data are not necessarily always useful for taxonomy, the current work showed the value of the ITS sequences as primary barcodes to help reveal interesting ecological phenomena.

Another matter that cannot be emphasized often enough is our strong recommendation not to discard new strains of fungi and other organisms after environmental or even physiological studies. If they end up in the global biodiversity repositories, other researchers may be able to make good use of them later on.

## Supplementary Information

The online version contains supplementary material available at XX

## Acknowledgements

The authors a wish to thank Aileen Gollasch for recording the HRESIMS spectra and Lucile Wendt for generating some of the DNA sequence data. We also thank Hans-Otto Baral for checking the “*Lachnum papyraceum*” specimen and revising the taxonomy.

## Author Contribution

The study including cultivation, extraction, data acquisition and analysis of the chromatogram was conducted by Özge Demir. The phylogeny was constructed by Yanpeng Chen based on rawdata generated by Lucile Wendt and curated by Christopher Lambert. The identity of the detected compounds was confirmed by Anja Schüffler, based on comparison with authentic standards. The first draft of the manuscript was written by and revised by F. Surup and M. Stadler. All authors read and approved the final version.

## Declarations

### Ethics approval

Not applicable.

### Consent to participate

Not applicable.

## Consent for publication

Not applicable.

## Footnotes

1 https://www.wipo.int/treaties/en/registration/budapest/

2 H.O.Baral (pers. comm.) stated that according to his experience the anamorphs of *Lachnum* are unknown, and this is not surprising, since none of the species seems to have been cultured and all are only known from teleomorphic descriptions.

3 https://www.inaturalist.org/

