## Supplementary Information for "Elucidation of the life cycle of a saprotrophic inoperculate discomycete that is associated with pinesap, using a polyphasic taxonomic approach^#^"

Özge Demir[a,b], Lucile Wendt[a,b], Yanpeng Chen[c], Christoper Lambert [a,b], Anja Schüffler[d], Frank Surup[a,b], Marc Stadler[a,b,*]

a. Department Microbial Drugs, Helmholtz Centre for Infection Research (HZI), German Centre for Infection Research (DZIF), Partner Site Hannover-Braunschweig, Inhoffenstrasse 7, 38124 Braunschweig, Germany
b. Institute of Microbiology, Technische Universität Braunschweig, Spielmannstraße 7, 38106 Braunschweig, Germany
c. School of Life Science and Technology, University of Electronic Science and Technology of China, Chengdu, 611731, People's Republic of China
d. Institut für Biotechnologie und Wirkstoff-Forschung gGmbH (IBWF), Hanns-Dieter-Hüsch-Weg 17, 55128 Mainz, Germany

### Alignment of the ITS sequences used in the phylogenetic study

>Lachnum_rachidicola_NBRC_114473

------------------------------------------------------------

---------------------------------------------------agtaaaagt

cgt---------------------------------------------------------

------------------------------------------------------------

------------------------------------------------------------

------------------------------------------------------------

-----------------------------------aacaaggtttccgt-aggtgaacct

gcggaaggatcattacag--agttcttgcccttc-------------------ggggtag
atctcccacccttgtgtatcattaataaatgttgctttggcgggccgtgagccta---gc
ttgcccgga-ttcgcgtccggcgtggcccgccagagaacccctaaactctgaatgt-tat
tgtcgtctgagtactatt-aaatagttaaaactttcaacaacggatctcttggttctggc
atcgatgaagaacgcagcgaaatgcgataagtaatgtgaattgcagaattcagtgaatca
tcgaatctttgaacgcacattgcgccccttggtattccgggggcatgcctgttcgagcg
tcattaataccaatctagcctggctaggtgttggact--tcgcgtctgggcgggtcttaa
aatcagtggcggtgctcttcagc-tctacgcgtagtaata-ctcctcgctatagggcctg
tggagacacttgccagcaaccccatatttttctaggttgacctcggatcaggtagggata
cccgctgaacttaagcata------------------------
>Lachnum_soppittii_FC__2160
------------------------------------------------------------
------------------------------------------------------------
------------------------------------------------------------
------------------------------------------------------------
------------------------------------------------------------
------------------------------------------------------------
--------------------------------------aaggtttccgt-aggtgaacct
gcggaaggatcattacag--agttcatgcccttc-------------------ggggtag
atctcccacccttgtgtatcattatagaatgttgctttggcgggctgcgtgccta---gc
gtgcccgga-tctcggtccggccc-gcccgccagaggacccttaaactctgaa-gt-tag
tgtcgtctgagtactatt-aaatagttaaaactttcaacaacggatctcttggttctggc
atcgatgaagaacgcagcgaaatgcgataagtaatgtgaattgcagaattcagtgaatca
tcgaatctttgaacgcacattgcgccccttggtattccgggggcatgcctgttcgagcg
tcatttataccaatctagcgtggctaggtgttgggct--tcgccgt-tggcgggccttaa
aattagtggcggtgctcttaggc-tctacgcgtagtaac--tttctcgctatagggtcct
aggagatgcttgccagcaacccc-aatttttctaggttgacc------------------
-------------------------------------------
>Lachnum_controversum_MFLU_18__1820
------------------------------------------------------------
------------------------------------------------------------
-gt---------------------------------------------------------

------------------------------------------------------------

------------------------------------------------------------

------------------------------------------------------------

-----------------------------------aacaaggtttccgt-aggtgaacct

gcggaaggatcattacag--agttcatgcccccc-------------------ggggtag

atctcccacccgtgtataccattatcgattgttgctttggcgggccgcgtgcccc---gc

acgcctggg-ctccggcccggcgc-gcccgccagaggacccccaaactctgaatgt-tag

tgtcgtctgagtactatt-aaatagttaaaactttcaacaacggatctcttggttctggc

atcgatgaagaacgcagcgaaatgcgataagtaatgtgaattgcagaattcagtgaatca

tcgaatctttgaacgcacattgcgccccctggcattccggggggcatgcctgttcgagcg

tcatttataccaatctcgcccagcgaggtgttgggct--ccgccgcctggcgggccctaa

aagcagtggcggtgcccttcggc-tctacgcgtagtaac--tttctcgctatagggtccg

gagggatgctggccagcaaccccaaatttttctaggttgacctcggatcaggtagggata

cccgctgaacttaagcatatctaa-------------------

>DSM10201

------------------------------------------------------------

-----------------------------------------------aggaagtaaaagt

cgt---------------------------------------------------------

------------------------------------------------------------

------------------------------------------------------------

------------------------------------------------------------

-----------------------------------aacaaggtttccgt-aggtgaacct

gcggaaggatcattacag--agttcatgcccttc-------------------ggggtag

atctcccacccttgtgtatcattatagaatgttgctttggcgggccgc--gcctt---gc

gcgcctaga-ttcgcgtctagcgt-gcccgccagaggacccctaaactctgaatgt-tag

tgtcgtctgagtactatt-aaatagttaaaactttcaacaacggatctcttggttctggc

atcgatgaagaacgcagcgaaatgcgataagtaatgtgaattgcagaattcagtgaatca

tcgaatctttgaacgcacattgcgccccctggtattccggggggcatgcctgttcgagcg

tcatttataccaatctagcctggctaggtgttgggct--tcgccgtctggcgggccttaa

aactagtggcggtgctcttaggc-tctacgcgtagtaat--tttctcgctatagggtcct

gggagatgcttgccagcaaccccaaatttttctaggttgacctcggatcaggtagggata

cccgctgaacttaagcatatcaata------------------

```
>Lachnum_virgineum_TNS__F__16583
---------------------------------cagcatccttccggggcgcttgccgaa
gcctatgcaaccctcacgggg------tccgcgactataaacac-----aggatgcaagt
cgcgatg--------gcggcaacataatcgaattg-cggggatctcctaaagccaagcag
taccaacaccgtccagaaatgtagggtgggccagcgttaatcccgctggaggttagaatc
tgctaggataccacaatggatgatccgcagcgaagcccctaacggcgccagcctacgggg
aacgttcacagactaagtggttgtgggtag-gc-tcgtcctgcttaagatatagtcgggc
cct----------------gtaggaaactacaggggctagccgttccgt-aggtgaacct
gcggaaggatcattacag--agttcatgcccttc-------------------ggggtag
atctcccacccttgtgtatcattatagaatgttgctttggcgggccgc--gcctc---gt
gcgcctaga-ttcgcgtctagcgt-gcccgccagaggatccctaaactctgaatgt-tag
tgtcgtctgagtactatt-aaatagttaaaactttcaacaacggatctcttggttctggc
atcgatgaagaacgcagcgaaatgcgataagtaatgtgaattgcagaattcagtgaatca
tcgaatctttgaacgcacattgcgccccctggtattccgggggggcatgcctgttcgagcg
tcatttataccaatctagcctggctaggtgttgggcg--ccgccgcctggcgggccttaa
aactagtggcggtgctctcaagc-tctacgcgtagtaat--cttctcgctacagggtctt
aggagatgctagccagcaatcccaaatttttctagg------------------------
-------------------------------------------
>ATCC36554
---------------------------------------ccttccgggtaacttgtggaa
gcctatgcaacctcaccaggg------tccgcgactataaatat-----aggatgcaagt
cgctatg--------gcggcaacatgatcgaattg-cggggatctcctaaagcctagcgg
taccaacacactgtagaaatgcagggtgggccagtgctaaccccactggaggtcagaatc
tgctaggatgctacaatggacgatccgcagcgaagcccctatcggcccaggcctacgggg
aacgttcacagactaggtggtcgtgggtag-gc-aagtcctgcttaagatatagtcgggc
gct----------------ctaggaaactacagcggtcaaccgttccgt-aggtgaacct
gcggaaggatcattacag--agttcatgcccttc-------------------ggggtag
atctcccacccttgtgtatcattatagaatgttgctttggcgggccgc--gcctc---gt
gcgcctagg-ttcgcgtctagcgt-gcccgccagaggacccctaaactctgaatgt-tag
tgtcgtctgagtactatt-aaatagttaaaactttcaacaacggatctcttggttctggc
atcgatgaagaacgcagcgaaatgcgataagtaatgtgaattgcagaattcagtgaatca
tcgaatctttgaacgcacattgcgccccctggtattccgggggggcatgcctgttcgagcg
```

tcatttataccaatctagcctggctaggtgttgggct--tcgccgtctggcgggccttaa

aactagtggcggtgctcttaggc-tctacgcgtagtaat--tttctcgctatagggtcct

gggagatgccggccagcaaccccaa-----------------------------------

-------------------------------------------

>Lachnum_abnorme_KUS__F52080

------------------------------------------------------------

------------------------------------------------------------

------------------------------------------------------------

------------------------------------------------------------

------------------------------------------------------------

------------------------------------------------------------

---------------------------------------------ccgtagggtgaacct

gcggaaggatcattacag--agttcatgcccttc-------------------ggggtag

acctcctacccttgtgtattataacaaattgttgccttggcgggctgcggctctg-----

ctgcccagg-attcgtcccggtac-gcccgccgaaggacccttaaactctgtttat-tag

tgtcgtctgagttttata-taatagttaaaactttcaacaacggatctcttggttctggc

atcgatgaagaacgcagcgaaatgcgataagtaatgtgaattgcagaattcagtgaatca

tcgaatctttgaacgcacattgcgccccttggtattccgaagggcatgcctgttcgagcg

tcattatgaccaatctagcctggctaggtattgggct--tcgcggttccgcgggccttaa

aaccagtggcggtgctctcaggc-tctacgcgtagtaac--tttctcgctttggagccct

gggggatgcttgccatcaacctc-aactttcttaggttgacctcggatcaggtagggata

cccgctgaacttaagcatatcaataagcggagga---------

>Trichopezizella_sp._KUS__F52478

------------------------------------------------------------

------------------------------------------------------------

------------------------------------------------------------

------------------------------------------------------------

------------------------------------------------------------

------------------------------------------------------------

--------------------------------------------------------gcct

gcggaaggatcattacag--agttcatgccctca-------------------cgggtag

atctcccacccttgtatatctat---cgatgttgctttagcgggccacaggccct---gc

ccgtaccgg-ccccggccggtcgt-gcccgctagagaacccc-aaactctgaatgt-tag
tgtcgtctgagtactata-aaatagttaaaactttcaacaacggatctcttggttctggc
atcgatgaagaacgcagcgaaatgcgataagtaatgtgaattgcagaattcagtgaatca
tcgaatctttgaacgcacattgcgccccttggtattccgaggggcatgcctgttcgagcg
tcatttaaaccaatctagctcagctaggtcttgggct--ccgccgtctggcgggccttaa
aaccagtggcggtactgccaggcttctaagcgtagtaaa--tttctcgctatagggccct
ggtggatgcttgccagcaacccctaatttttcaggttgacctcggatcaggtagggata
cccgctgaacttaagcatatcaaaaggcc--------------
>Trichopeziza_sulphurea_KUS__F52218
------------------------------------------------------------
------------------------------------------------------------
------------------------------------------------------------
------------------------------------------------------------
------------------------------------------------------------
------------------------------------------------------------
-------------------------------------------ttccgt-aggtgaacct
gcggaaggatcattacag--agttcatgccctca-------------------cgggtag
atctcccacccttgtgtattata-tcatatgttgctttggcgggt---------------
--------------------gcaa-gcccgccagaggaccc--aaactctgagt-t-tag
tgtcgtctgagtactata-aaatagttaaaactttcaacaacggatctcttggttctggc
atcgatgaagaacgcagcgaaatgcgataagtaatgtgaattgcagaattcagtgaatca
tcgaatctttgaacgcacattgcgccccttggtattccgaggggcatgcctgttcgagcg
tcattataaccaatccagcctggctgggtcttgggct--ccgccaaccggcgggccttaa
aatcagtggcggtgctgccaggc-tctaagcgtagtaaa--tacctcgctatagggtcct
ggtggaggcttgccaacaacccccaacttct-taggttgacctcggatcaggtagggata
cccgctgaacttaagcatatcaataagcggag-----------
>Lachnum_fuscescens_FC__2200
----------------------------------agtatccttcccggtaactttcggaa
gcctttgcaacctaatcaggg------tccgcgactataaatac-----cggatgcaagt
agcctaga-------gctgcaacatgatcgaattg-cggggacgtcctaaagccta-cga
taccaacgtttcagggaaacctgcagcgggct-gtgtaaac------agaaggtcataac
tcgtaggatg-tacaatggataatccgcagcgaagcccctaacggtgtaagcctacgggg

aacgttcacagactaagtggttgtgggtagagc-acgctctgcttaagatatagtcggac

gct----------------gtgggaaactgcagtgactaatcgttccgt-aggtgaacct

gcggaaggatcattacag--agttcttgccctaa-------------------cgggtag

aattcccacccttgtgtaacataattatctgttgctttggcgggtcgcgagttta-----

------------cttgccctgtgt-acccgccagagaaccaatcaactctgaatat-tag

tgtcgtctgagtactata-taatagttaaaactttcaacaacggatctcttggttctggc

atcgatgaagaacgcagcgaaatgcgataagtaatgtgaattgcagaattcagtgaatca

tcgaatctttgaacgcacattgcgccccttggtattccgggggcatgcctgttcgagcg

tcattataaccaatccagctt-gctgggtcttgggctatcgcctccctggcgggccttaa

aaacagtggcggtgctctcaagc-tctacgcgtagtaatt-attctcgctattgggtctt

gagggatgcttgccaacaacccccaacttct-taggttgacctcggat------------

-------------------------------------------

>Brunnipila_fuscescens_KUS__F52031

------------------------------------------------------------

------------------------------------------------------------

------------------------------------------------------------

------------------------------------------------------------

------------------------------------------------------------

------------------------------------------------------------

-------------------------------------------ttccgtagggtgaacct

gcggaaggatcattacag--agttcatgccctaa-------------------cgggtag

atctcccacccttgtgtatcataattatctgttgctttggcggggcgcgagttcg-----

------------ctcgccctgtgt-acccgccagaggatcaccaaactctgaatgt-tag

tgtcgtctgagtactata-aaatagttaaaactttcaacaacggatctcttggttctggc

atcgatgaagaacgcagcgaaatgcgataagtaatgtgaattgcagaattcagtgaatca

tcgaatctttgaacgcacattgcgccccttggtattccgggggcatgcctgttcgagcg

tcattataaccaatccagctt-gctgggtcttgggctatcgcctccctggcgggccttaa

aaacagtggcggtgctctcaagc-tctacgcgtagtaattatttctcgctatagggtctt

gagggatgcttgccatcaacccccaatttttctaggttgacctcggatcaggtagggata

cccgctgaacttaagcatatcaataagcggag-----------

>Proliferodiscus_sp._KUS__F52660

------------------------------------------------------------

------------------------------------------------------------
------------------------------------------------------------
------------------------------------------------------------
------------------------------------------------------------
------------------------------------------------------------
------------------------------------------------tatagggnacct
gcggaaggatcattacag--agtccctgcccctct------------------ggggtag
acctcccacccgtgtgtattctaatgaattgttgctttggcgggccgcgagcgtg---cc
tcgcccgggagcctctcccggtgt-gcccgccagaggaaacccaaactctgaatat-tag
tgtcgtctgagtactacataaatcgttaaaactttcaacaacggatctcttggttctggc
atcgatgaagaacgcagcgaaatgcgataagtaatgtgaattgcagaattcagtgaatca
tcgaatctttgaacgcacattgcgccccttggtattccgggggggcatgcttgttcgagcg
tcattatgacccctccagcctagctgggtgttgggcc--gcgcctccgggcgggtcttaa
aactagtggcggtgctttcaggc-tctacgcgtagtaact-tttctcgctatagggtcct
gggagatgctagccagcaacctcaaactttttcaggttgacctcgaatcaagtagggata
cccgctgaacttaagcatatcaataagcggaggaagatcatta
>Proliferodiscus_sp._TNS__F17436
------------------------------------------------------------
------------------------------------------------------------
------------------------------------------------------------
------------------------------------------------------------
------------------------------------------------------------
------------------------------------------------------------
----------------------------------------------------gtgaacct
gcggaaggatcattacag--agtccctgcccctct------------------ggggtag
acctcccacccgtgtgtattctaatgaattgttgctttggcgggccgcgagcgtg---cc
tcgcccgggagcctctcccggtgt-gcccgccagaggaaacccaaactctgaatat-tag
tgtcgtctgagtactacataaatcgttaaaactttcaacaacggatctcttggttctggc
atcgatgaagaacgcagcgaaatgcgataagtaatgtgaattgcagaattcagtgaatca
tcgaatctttgaacgcacattgcgccccttggtattccgggggggcatgcttgttcgagcg
tcattatgacccctccagcctagctgggtgttgggcc--gcgcctccgggcgggtcttaa
aactagtggcggtgctttcaggc-tctacgcgtagtaact-tttctcgctatagggtcct

gggagatgctagccagcaacctcaaactttttcaggttgacctcgaatcaagtagggata

cccgctgaacttaagcatatcaataa----------------

>Lachnum_palmae_TNS_F__24600

------------------------------------------------------------

------------------------------------------------------------

------------------------------------------------------------

------------------------------------------------------------

------------------------------------------------------------

------------------------------------------------------------

---------------------------------------atccttccgt-aggggaacct

gcggaaggatcattacag--agttcgtgcccttt---------------------gggta

gttctccacccttgtgtatttaa---ccacgttgcttcggcgggtt--------------

-----------------------c-gcccgccaaaggccc---aaactctatatatcttg

tatcccctgagtcttata-taatatttaaaactttcaacaacggatctcttggttctggc

atcgatgaagaacgcagcgaaatgcgataagtaatgtgaattgcagaattcagtgaatca

tcgaatctttgaacgcacattgcgccccttggtattccgaggggcatgcctgttcgagcg

tcattatacccaatctagctt-gctaggtcttggacc----cgcggttcgcgggtcttaa

aatcagcggcggtgcccc-tggc-tctatgcgtagtaa---tttctcgctccaga-cccc

agggaccacctgccaaaa------aacttctttaggttgacctcggatcaggtagggata

cccgctgaacttaagcatatcaataagcggaggaa--------

>Arachnopeziza_aurelia_TNS__F11211

ggatttagtgttttactattggtattacgctatgatccttccacagtgacacttatagaa

gcctttgcagcctggcaacaggtgctccatgcgacagtaaataagttggacgatgcaagt

cataagagtaatcttgtggcgacatgatcgaattgacggggacatcctaaagcttg-cca

caccaacttttcggggaaactcgaggagggcctatgttaactgcataggagggtaaaaga

gggcaagatattacaatggacaatccgcagcgaagaccctaagtgcact-gcatatgggt

aacgttcacagactaagtggttgtgggtggcacgatgtgctgcttaagatatagtcgggc

ctttagtgaaagctaaggggtaagtcacaaagacgattaaccgttccgt-aggtgaacct

gcggaaggatcattaaaa--agtttcggtctataaggcgtaaaaccctctatagaccgtg

aacccccacccttgtgtattat--ttatttgttgctttggcaggccgcgagctttacagc

gagcaccggcttcggctggagagt-gcctgccagaggaccc---aactctgtattt-tag

tgacatctgagtactata-taatagttaaaactttcaacaacggatctcttggttctggc

atcgatgaagaacgcagcgaaatgcgataagtaatgtgaattgcagaattcagtgaatca
tcgaatctttgaacgcacattgcgccctctggtattccgggggcatgcctgttcgagcg
tcattatgaccaatcacgcaag---tggtattggggc--ttgccatacggcatcccttaa
acgcagtggcagtgctat-aggc-tctcagcgcagtaat--tatttcgctctcgagtcct
-agacccacccgccaaaa-ccccaacttctt---aggtgacctcgg--------------
-------------------------------------------
>Arachnopeziza_aurata_TNS__F11212
------------------------------------------------------------
-----------------------------------------------aggaagtaaaagt
cgt---------------------------------------------------------
------------------------------------------------------------
------------------------------------------------------------
------------------------------------------------------------
-----------------------------------aacaaggtttccgt-aggtgaacct
gcggaaggatcattagaatgagattcggtccgagggcttaaaaacctt---agggccgtc
aacctccacccttgtgtatcat--tcaattgttgctttggtgggccgcgacccgcacggg
gagcaccggcttcagctggagagt-gcctgccagagaacccc--aactctgtatt--ttg
taacatctgagtactata-caatagttaaaactttcaacaacggatctcttggttctggc
atcgatgaagaacgcagcgaaatgcgataagtaatgtgaattgcagaattcagtgaatca
tcgaatctttgaacgcacattgcgccctctggtattccgggggcatgcctgttcgagcg
tcattatgaccaatcacgcaag---tggtattggggc--ttgccgt-cggcatcccttaa
aatcagtggcagtgctat-aggc-tctcagcgcagtaat--tatttcgctcttgagtcct
-agacccacccgccaaaa-ccccaactattttaaggttgacctcggatcaggtagggata
cccgctgaacttaagcatatc----------------------
>Lachnum_rachidicola_NBRC_114473
tagcatatcaataagcggaggaaaagaaaccaacagggattacctcagtaacggcgagtg
aagcggtaacagctcaaatttgaaatctggctctttcagggtccgagttgtaatttgtag
aagatgctttgggtgtggtccgggtctaagttccttggaacaggacgtcatagagggtga
gaatcccgtatgtgactcggtaccttcgcccgtgtaaagctctttcgacgagtcgagttg
tttgggaatgcagctcaaaatgggtggtaaatttcatctaaagctaaatattggccagag
accgatagcgcacaagtagagtgatcgaaagatgaaaagcactttggaaagagagttaaa
cagtacgtgaaattgttgaaagggaagcgcttgcaaccagacttagcggcggctgatcat

ccggggttctccccggtgcactcggtcgtctctaggccagcatcggtttgggtggtggga
taaaggccttggtaatgtagcttctctcggggagtgttatagacctcggtgcaatgccgc
ctacccggaccgaggaccgcgcttcggctaggatgctggcgtaatggttgtaagcgaccc
gtcttgaaacacggaccaaggagtctaacatctatgcgagtatttgggtgtc-aaaccca
tatgcgtaatgaaagtgaacggaggtgagaaccctt-aagggtgcatcatcgaccgatcc
tgatgtcttcggatggatttgagtaagagcatagctgttgggacccg-aaagatggtgaa
ctatgcctaaatagggtgaagccagaggaaactctggtggaggctcgcagcggttctgac
gtgcaaatcgatcgtcaaatttgggtataggggcgaaagactaatcgaaccat-------
------------------------------------------------------------
------------------------------------------------------------
------------------------------------------------------------
------------------------------------------------------------
------------------------------------ctagtagctgg-------------
------------------------------------------------------------
------------------------------------------------------------
------------------------------------------------------------
------------------------------------------------------------
------------------------------------------------------------
------------------------------------------------------------
------------------------------------------------------------
---------------------------------------------------------
>Lachnum_virgineum_TNS__F__16583
---------------------------------------------cagtaacggcgagtg
aagcggtaacagctcaaatttgaaatctggctctttcagggtccgagttgtaatttgtag
aagatgctttgggtgtggccctggtctaagttccttggaacaggacgtcatagagggtga
gaatcccgtatgtgactaggtgctttcgcccgtgtaaagctctttcgacgagtcgagttg
tttgggaatgcagctcaaaatgggtggtaaatttcatctaaagctaaatattggccagag
accgatagcgcacaagtagagtgatcgaaagatgaaaagcactttggaaagagagttaaa
cagtacgtgaaattgttgaaagggaagcgcttgcaaccagacttagttgcggccgatcat
ctagggttctccctggtgcactcggtcgtatctaggccagcatcggtttgggtggtggga
taaaggccttgggaatgtagcttctttcggggagtgttatagccctcggtgcaatgccgc
ctacctggaccgaggaccgcgcttcggct-------------------------------

------------------------------------------------------------
------------------------------------------------------------
------------------------------------------------------------
------------------------------------------------------------
------------------------------------------------------------
------------------------------------------------------------
------------------------------------------------------------
------------------------------------------------------------
------------------------------------------------------------
------------------------------------------------------------
------------------------------------------------------------
------------------------------------------------------------
------------------------------------------------------------
------------------------------------------------------------
------------------------------------------------------------
------------------------------------------------------------
------------------------------------------------------------
---------------------------------------------------------
>DSM10201
---------------------------------------------------cggcgagtg
aagcggtaacagctcaaatttgaaatctggctcttttagggtccgagttgtaatttgtag
aagatgctttgggtgtggccctggtctaagttccttggaacaggacgtcatagagggtga
gaatcccgtatgtgactaggtgtcttcgcccgtgtaaagctctttcgacgagtcgagttg
tttgggaatgcagctcaaaatgggtggtaaatttcatctaaagctaaatattggccagag
accgatagcgcacaagtagagtgatcgaaagatgaaaagcactttggaaagagagttaaa
cagtacgtgaaattgttgaaagggaagcgcttgcaaccagacttagttgcggccgatcat
ctagggttctccctggtgcactcggtcgtatctaggccagcatcggtttgggtggtggga
taaaggccttgggaatgtagcttctttcggggagtgttatagccctcggtgcaatgccgc
ctacctggaccgaggaccgcgcttcggctaggatgctggcgtaatggttgtaagcgaccc
gtcttgaaacacggaccaaggagtctaacatctatgcgagtatttgggtgtt-aaaccca
tatgcgtaatgaaagtgaacggaggtgagaacccctt-aagggtgcatcatcgaccgatcc
tgatgtcttcggatggatttgagtaagagcatagctgttgggacccg-aaagatggtgaa

ctatgcctaaatagggtgaagccagaggaaactctggtggaggctcgcagcggttctgac

gtgcaaatcgatcgtcaaatttgggtataggggcgaaagactaatcgaaccat-------

------------------------------------------------------------

------------------------------------------------------------

------------------------------------------------------------

------------------------------------------------------------

------------------------------------ctagtagctggttcctgccgaag-

---------------tttccctcaggatagcagtgttgaatt------------------

----cagttttatgaggtaaagcgaatgattagaggccttggggttgaaacaaccttaac

ctattctcaaactttaaatatgtaagaagtccttgttacttaattgaacgtggacattcg

aatgtaccaacactagtgggccatttttggtaagcagaactggcgatgcgggatgaaccg

aacgtgaagttaaggtgccggaatat----acgctcatcagacaccacaaaaggtgttag

ttcatctagacagcaggacggtggccatggaagtcggaatccgctaaggaatgtgtaaca

actcacctgccgaatgaactagccctgaaaatggatggcgcttaagcgtattacccatac

ttcaccgccagggtagaaacgatgccctggcgagtaggc------------------

>ATCC36554

------------------------------------------------------------

--gcggtaacagctcaaatttgaaatctggctctttcagggtccgagttgtaatttgtag

aagatgctttgggtgtggccctggtctaagttccttggaacaggacgtcatagagggtga

gaatcccgtatgtgactaggtgctttcgcccgtgtaaagctctttcgacgagtcgagttg

tttgggaatgcagctcaaaatgggtggtaaatttcatctaaagctaaatattggccagag

accgatagcgcacaagtagagtgatcgaaagatgaaaagcactttggaaagagagttaaa

cagtacgtgaaattgttgaaagggaagcgcttgcaaccagacttagttgcggccgatcat

ctagggttctccctggtgcactcggtcgtatctaggccagcatcggtttgggtggtggga

taaaggccttgggaatgtagcttctttcggggagtgttatagccctcggtgcaatgccgc

ctacctggaccgaggaccgcgcttcggctaggatgctggcgtaatggttgtaagcgaccc

gtcttgaaacacggaccaaggagtctaacatctatgcgagtatttgggtgtt-aaaccca

tatgcgtaatgaaagtgaacggaggtgagaaccctt-aagggtgcatcatcgaccgatcc

tgatgtcttcggatggatttgagtaagagcatagctgttgggacccg-aaagatggtgaa

ctatgcctaaatagggtgaagccagaggaaactctggtggaggctcgcagcggttctgac

gtgcaaatcgatcgtcaaatttgggtataggggcgaaagactaatcgaaccat-------

------------------------------------------------------------

------------------------------------------------------------
------------------------------------------------------------
------------------------------------------------------------
------------------------------------ctagtagctggttcctgccgaag-
---------------tttccctcaggatagcagtgttgaatt------------------
----cagttttatgaggtaaagcgaatgattagaggccttggggttgaaacaaccttaac
ctattctcaaactttaaatatgtaagaagtccttgttacttaattgaacgtggacattcg
aatgtaccaacactagtgggccatttttggtaagcagaactggcgatgcgggatgaaccg
aacgtgaagttaaggtgccggaatat----acgctcatcagacaccacaaaaggtgttag
ttcatctagacagcaggacggtggccatggaagtcggaatccgctaaggaatgtgtaaca
actcacctgccgaatgaactagccctgaaaatggatggcgcttaagcgtattacccatac
ttcaccgccagggtagaaacgatgccctggcgagtaggcaggcgtgggggtca----
>Lachnum_soppittii_FC__2160
--------------------------------------attacctcagtaacggcgagtg
aagcggtaacagctcaaatttgaaatctggctctttcagggtccgagttgtaatttgtag
aagatgctttgggtgtggcccgggtctaagttccttggaacaggacgtcatagagggtga
gaatcccgtatgtgacccggcgctttcgcccgtgtaaagctctttcgacgagtcgagttg
tttgggaatgcagctcaaaatgggtggtaaatttcatctaaagctaaatattggccagag
accgatagcgcacaagtagagtgatcgaaagatgaaaagcactttggaaagagagttaaa
cagtacgtgaaattgttgaaagggaagcgcttgcaaccagacttaggcgcggccgatcat
ccagggttctccctggtgcactcggtcgtctctaggccagcatcggtttgggtggtggga
taaaggccttgggaatgtagcttctttcggggagtgttatagccctcggtgcaatgccgc
ctacctggaccgaggaccgcgcttcggctaggatg-------------------------
------------------------------------------------------------
------------------------------------------------------------
------------------------------------------------------------
------------------------------------------------------------
------------------------------------------------------------
------------------------------------------------------------
------------------------------------------------------------
------------------------------------------------------------
------------------------------------------------------------

------------------------------------------------------------
------------------------------------------------------------
------------------------------------------------------------
------------------------------------------------------------
------------------------------------------------------------
------------------------------------------------------------
------------------------------------------------------------
------------------------------------------------------------
---------------------------------------------------------
>Lachnum_controversum_MFLU_18__1820
-----------aatagcggagaaaagaaaccaacagggattacctcagtaacggcgagtg
aagcggtaacagctcaaatttgaaatctggctctttcagggtccgagttgtaatttgtag
aagatgctttgggtgtggcccgggtctaagttccttggaacaggacgtcatagagggtga
gaatcccgtatgtgactcggcgccttcgcccgtgtaaagctctttcgacgagtcgagttg
tttgggaatgcagctcaaaatgggtggtaaatttcatctaaagctaaatattggccggag
accgatagcgcacaagtagagtgatcgaaagatgaaaagcactttggaaagagagttaaa
cagtacgtgaaattgttgaaagggaagcgcttgcaaccagacttagttgctgtcgatcat
ctcgggttctcccgggtgcactcggcagtgtctaggccagcatcggtttgggtggtggga
taaaggccttgggaatgtagcttctttcggggagtgttatagccctcggtgcaatgccgc
ctacccggaccgaggaccgcgcttcggctaggatgctggcgtaatggttgtaagcgaccc
gtcttgaaacacggaccaaggagtctaacatctatgcgagtatttgggtgtc-aaaccca
tatgcgtaatgaaagtgaacggaggtgagaaccctt-aagggtgcatcatcgaccgatcc
tgatgtcttcggatggatttgagtaagagcatagctgttgggacccg-aaagatggtgaa
ctatgcctaaatagggtgaagccagaggaaactctggtggaggctcgcagcggttctgac
gtgcaaatcgatcgtcaaatttgggtatagggggcgaaagactaatcgaac-at-------
------------------------------------------------------------
------------------------------------------------------------
------------------------------------------------------------
------------------------------------------------------------
------------------------------------ctag-agctggtcc----------
------------------------------------------------------------
------------------------------------------------------------

------------------------------------------------------------
------------------------------------------------------------
------------------------------------------------------------
------------------------------------------------------------
------------------------------------------------------------
---------------------------------------------------------
>Trichopeziza_sulphurea_KUS__F52218
------------------------------------------------------------
--------------caaatttgaaatctggccctttcagggtccgagttgtaatttgtag
aagatgctttgggtgtggctccggtctaagttccttggaacaggacgtcatagagggtga
gaatcccgtatgtgaccggtggctttcgcccgtgtaaagctctttcgacgagtcgagttg
tttgggaatgcagctcaaaatgggtggtatatttcatctaaagctaaatattggccagag
accgatagcgcacaagtagagtgatcgaaagatgaaaagcactttggaaagagagttaaa
cagtacgtgaaattgttgaaagggaagcgcttgcaaccagactcgcgcgccgttgatcat
ccggggttctccccggtgcactcggcggcgctcgggccagcatcagtttgggtggtggga
taaaggccttgggaatgtagctcccctcggggagtgttatagccctcggtgcaatgccac
ctacccggactgaggaccgcgcttcggctaggatgctggcgtaatggttgtaagcgaccc
gtcttgaaacacggaccaaggagtctaacatctatgcgagtgtttgggtgtc-aaaccca
tacgcgtaatgaaagtgaacggaggtgagaacccttt-aagggtgcatcatcgaccgatcc
tgatgtcttcggatggatttgagtaagagcatagctgttgggacccg-aaagatggtgaa
ctatgcctaaatagggtgaagccagaggaaactctggtggaggctcgcagcggttctgac
gtgcaaatcgatcgtcaaatttgggtataggggcgaaagactaatcgaaccat-------
------------------------------------------------------------
------------------------------------------------------------
------------------------------------------------------------
------------------------------------------------------------
------------------------------------ctagtagctggttcctgccgaag-
---------------tttccctcaggatagcagtgttgaatt------------------
----cagttttatgaggtaaagcgaatgattagaggccttggggatgcaacatccttaac
ctattctcaaactttaaatatgtaagaagtccttgttacttaattgaacgtggacattcg
aatgtaccaacactagtgggccatttttggtaagcagaactggcgatgcgggatgaaccg
aacgtgaagttaaggtgccggaatat----acgctcatcagacaccacaaaaggtgttag

ttcatctagacagcaggacggtggccatggaagtcggaatccgctaaggagtgtgtaaca

actcacctgccgaatgaactagccctgaaaatggatggcgcttaagcgtattacccatac

ttcaccgcc---------------------------------------------------

>Brunnipila_fuscescens_KUS__F52031

------------------------------------------------taacggcgagtg

aagcggtaacagctcaaatttgaaatctggctctctcagggtccgagttgtaatttgtag

aagatgctttgagtgtggctctagtctaagttccttggaacaggacgtcatagagggtga

gaatcccgtatgtgattaggcgccttcgctcgtgtaaagctctttcgacgagtcgagttg

tttgggaatgcagctcaaaatgggtggtaaatttcatctaaagctaaatattggccagag

accgatagcgcacaagtagagtgatcgaaagatgaaaagcactttggaaagagagttaaa

cagtacgtgaaattgttgaaagggaagcgcttgcaaccagactcgcatgccgtcgatcat

cctgtgttctcactggtgcactcggcggccttcgggccagcatcggtttgggtggtggga

taaaggccttgggaatgtagctcctctcggggagtgttatagccctcggtgcaatgccgc

ctacctggaccgaggaccgcgcttcggctaggatgctggcgtaatggttgtaagcgaccc

gtcttgaaacacggaccaaggagtctaacatctatgcgagtatttgggtgtc-aaaccca

tatgcgtaatgaaagtgaacggaggtgagaaccctt-aagggtgcatcatcgaccgatcc

tgatgtcttcggatggatttgagtaagagcatagctgttgggacccg-aaagatggtgaa

ctatgcctaaatagggtgaagccagaggaaactctggtggaggctcgcagcggttctgac

gtgcaaatcgatcgtcaaatttgggtataggggcgaaagactaatcgaaccat-------

------------------------------------------------------------

------------------------------------------------------------

------------------------------------------------------------

------------------------------------------------------------

------------------------------------ctagtagctggttcctgccgaag-

---------------tttccctcaggatagcagtgttgaatt------------------

----cagttttatgaggtaaagcgaatgattagaggccttggggttgaaacaaccttaac

ctattctcaaactttaaatatgtaagaagtccttgttacttaattgaacgtggacattcg

aatgtaccaacactagtgggccatttttggtaagcagaactggcgatgcgggatgaaccg

aacgtgaagttaaggtgccggaatat----acgctcatcagacaccacaaaaggtgttag

ttcatctagacagcaggacggtggccatggaagtcggaatccgctaaggaatgtgtaaca

actcacctgccgaatgaactagccctgaaaatggatggcgcttaagcgtattacccatac

ttcaccgccagggtagaaacgatgccctggcgagtaggcaggcgtggagg-------

>Trichopezizella_sp._KUS__F52478

------------------------------------------------------------

------------------tttgaaatctggctcttttagggtccgagttgtaatttgtag

aagatgctttgggtgtggctccggtctaagttccttggaacaggacgtcatagagggtga

gaatcccgtatgtgactggttgccttcgcccgtgtaaagctctttcgacgagtcgagttg

tttgggaatgcagctcaaaatgggtggtatatttcatctaaagctaaatattggccagag

accgatagcgcacaagtagagtgatcgaaagatgaaaagcactttggaaagagagttaaa

cagtacgtgaaattgttgaaagggaagcgcttgcaaccagactcgcgcgctgttgatcat

ccggtgttctcaccggtgcactcagcagtgctcgggccagcatcagtttgggtggtggga

taaaggccttgggaatgtagcttctttcggggagtgttatagccctcggtgcaatgccgc

ctacctggactgaggaccgcgcttcggctaggatgctggcgtaatggttgtaagcgaccc

gtcttgaaacacggaccaaggagtctaacatctatgcgagtgtttgggtgtc-aaaccca

tacgcgtaatgaaagtgaacggaggtgagaacccttaaagggtgcatcatcgaccgatcc

tgatgtcttcggatggatttgagtaagagcatagctgttgggacccg-aaagatggtgaa

ctatgcctaaatagggtgaagccagaggaaactctggtgga-gctcgcagcggttctgac

gtgcaaatcgatcgtcaaatttgggtataggggcgaaagactaatcg-accat-------

------------------------------------------------------------

------------------------------------------------------------

------------------------------------------------------------

------------------------------------------------------------

------------------------------------ctagtagctgt-------------

------------------------------------------------------------

------------------------------------------------------------

------------------------------------------------------------

------------------------------------------------------------

------------------------------------------------------------

------------------------------------------------------------

------------------------------------------------------------

---------------------------------------------------------

>Proliferodiscus_sp._KUS__F52660

------------------------------------------------------------

--------------caaatttgaaatctggctctttcagggtccgagttgtaatttgtag aagatgctttgggtgtggccccggtctaagttccttggaacaggacgtcatagagggtga

gaatcccgtatgtgactgggtgctttcgcccgtgtaaagctctttcgacgagtcgagttg

tttgggaatgcagctcaaaatgggtggtaaatttcatctaaagctaaatattggccagag

accgatagcgcacaagtagagtgatcgaaagatgaaaagcactttggaaagagagttaaa

cagtacgtgaaattgttgaaagggaagcgcttgcaaccagactcgcatgccgctgatcat

ccggggttctccccggtgcactcggtggtctacgggccagcatcggtttgggtggcggga

taaaggctgtgggaatgtagcttctctcggggagtgttatagcccacggtgcaatgccgc

ctacctggaccgaggaccgcgcttcggctaggatgctggcgtaatggttgtaagcgaccc

gtcttgaaacacggaccaaggagtctaacatctatgcgagtatttgggtgtt-aaaccca

tatgcgtaatgaaagtgaacggaggtaagaaccctt-tagggtgcattatcgaccgatcc

tgatgtcttcggatggatttgagtaagagcatagctgttgggacccgaaaagatggtgaa

ctatgcctaaatagggtgaagccagaggaaactctggtggaggctcgcagcggttctgac

gtgcaaatcgatcgtcaaatttgggtataggggcgaaagactaatcgaaccat-------

------------------------------------------------------------

------------------------------------------------------------

------------------------------------------------------------

------------------------------------------------------------

------------------------------------ctagtagtgtc-------------

------------------------------------------------------------

------------------------------------------------------------

------------------------------------------------------------

------------------------------------------------------------

------------------------------------------------------------

------------------------------------------------------------

------------------------------------------------------------

---------------------------------------------------------

>Proliferodiscus_sp._TNS__F17436

-------------agcggaggaaaagaaaccaacagggattacctcagtaacggcgagtg

aagcggtaacagctcaaatttgaaatctggctctttcagggtccgagttgtaatttgtag

aagatgctttgggtgtggccccggtctaagttccttggaacaggacgtcatagagggtga

gaatcccgtatgtgactgggtgctttcgcccgtgtaaagctctttcgacgagtcgagttg

tttgggaatgcagctcaaaatgggtggtaaatttcatctaaagctaaatattggccagag

accgatagcgcacaagtagagtgatcgaaagatgaaaagcactttggaaagagagttaaa
cagtacgtgaaattgttgaaagggaagcgcttgcaaccagactcgcatgccgctgatcat
ccggggttctccccggtgcactcggtggtctacgggccagcatcggtttgggtggcggga
taaaggctgtgggaatgtagcttctctcggggagtgttatagcccacggtgcaatgccgc
ctacctggaccgaggaccgcgcttcggctaggatgctggcgtaatggttgtaagcgaccc
gtcttgaaa---------------------------------------------------
------------------------------------------------------------
------------------------------------------------------------
------------------------------------------------------------
------------------------------------------------------------
------------------------------------------------------------
------------------------------------------------------------
------------------------------------------------------------
------------------------------------------------------------
------------------------------------------------------------
------------------------------------------------------------
------------------------------------------------------------
------------------------------------------------------------
------------------------------------------------------------
------------------------------------------------------------
------------------------------------------------------------
------------------------------------------------------------
---------------------------------------------------------
>Arachnopeziza_aurelia_TNS__F11211
-------------agcggaggaaaagaaaccaacagggattacctcagtaacggcgagtg
aagcggtaacagctcaaatttgaaatctggctctc---gggcccgagttgtaatttgtag
aagatgctttgggtgtggctccggtctaagttccttggaacaggacgtcatagagggtga
gaatcccgtatgtgactggtcgccttcgcccgtgtaaagctctttcgacgagtcgagttg
tttgggaatgcagctcaaaatgggtggtatatttcatctaaagctaaatattggccagag
accgatagcgcacaagtagagtgatcgaaagatgaaaagcactttggaaagagagttaaa
cagtacgtgaaattgttgaaagggaagcgcttgcaaccagacttgcaggctgttgatcat
ccggggttctccccggtgcactcggcagttttcaggccagcatcggtttgagtggtggga

taaaggccttgggaatgtggctctcttcggggagtgttatagcccaaggtgcaatgccgc

ctacttggaccgaggaccgcgcttcggctaggatgctggcgtaatggttgtaagcgaccc

gtcttg------------------------------------------------------

------------------------------------------------------------

------------------------------------------------------------

------------------------------------------------------------

------------------------------------------------------------

------------------------------------------------------------

------------------------------------------------------------

------------------------------------------------------------

------------------------------------------------------------

------------------------------------------------------------

------------------------------------------------------------

------------------------------------------------------------

------------------------------------------------------------

------------------------------------------------------------

------------------------------------------------------------

------------------------------------------------------------

------------------------------------------------------------

---------------------------------------------------------

>Arachnopeziza_aurata_TNS__F11212

-------------agcggaggaaaagaaaccaacagggattacctcagtaacggcgagtg

aagcggtaacagctcaaatttgaaatctggctctc---gggcccgagttgtaatttgtag

aagatgctttgggtgtggctccggtctaagttccttggaacaggacgtcatagagggtga

gaatcccgtacgtgactggtggccttcgctcgtgtaaagctctttcgacgagtcgagttg

tttgggaatgcagctcaaaatgggtggtatatttcatctaaagctaaatattggccagag

accgatagcgcacaagtagagtgatcgaaagatgaaaagcactttggaaagagagttaaa

cagtacgtgaaattgttgaaagggaagcgcttgcaaccagacttgcaggctgttgatcat

ccggggttctccccggtgcactcggcagttttcaggccagcatcggtttgggtggtggga

taaaggccttgggaatgtggctctcttcggggagtgttatagcccaaggtgcaatgccgc

ctacctggaccgaggaccgcgcttcggctaggatgctggcgtaatggttgtaagcgaccc

gtcttg------------------------------------------------------

------------------------------------------------------------

------------------------------------------------------------

------------------------------------------------------------

------------------------------------------------------------

------------------------------------------------------------

------------------------------------------------------------

------------------------------------------------------------

------------------------------------------------------------

------------------------------------------------------------

------------------------------------------------------------

------------------------------------------------------------

------------------------------------------------------------

------------------------------------------------------------

------------------------------------------------------------

------------------------------------------------------------

------------------------------------------------------------

---------------------------------------------------------

>Lachnum_abnorme_KUS__F52080

--------------------------------acagggattacctcagtaacggcgagtg

aagcggtaacagctcaaatttgaaatctggctctttcagggtccgagttgtaatttgtag

aagatgctttgggcgtggctccagtctaagttccttggaacaggacgtcatagagggtga

gaatcccgtatgtgattggtggctttcgcccgtgtaaagctctttcgacgagtcgagttg

tttgggaatgcagctcaaaatgggtggtaaatttcatctaaagctaaatattggccagag

accgatagcgcacaagtagagtgatcgaaagatgaaaagcactttggaaagagagttaaa

cagtacgtgaaattgttgaaagggaagcgttggcaaccagactcgtatgccgctaatcat

ccggggttctccccggtgcacttggtggttttcgggccagcatcggtttcggtggtggga

taaaggccttgggaatgtagcttctctcggggagtgttatagccctcggtgcaatgccgc

ctactgggaccgaggaccgcgcttcggctaggatgctggcgtaatggttgtcaacggccc

gtcttgaaacacggaccaaggagtctaacatctatgcgagtatttgggtgtc-aaaccca

tatgcgtaatgaaagtgaacggaggtaagagccctt-aagggtgcattatcgaccgatcc

tgatgtcttcggatggatttgagtaagagcatagctgttgggacccg-aaagatggtgaa

ctatgcctaaatagggtgaagccagaggaaactctggtggaggctcgcagcggttctgac

gtgcaaatcgatcgtcaaattcgggcataggggcgaaagactaatcgaaccattggaata
cctactaggtggttaagaggcgtaagcctagtcctgcacagggcaacattgtcaaattgt
tcggggacctcccgcacttctcagctaccgcagcctggccgaaaggcgggcgcgcaccag
ggtaacgccctcggggatggtaagaacgctgaaaaggggacgatccgcagcttcttccta
cgggcttcgcctacggagggagttcacagactcgatggcagtgggcctctgggcttaaga
tagagtcgaaccaccgggcaaccggatggagcaatgctagtagctggttcctgccgaag-
---------------tttccctcaggatagcagtgttgtttt------------------
----cagttttatgaggtaaagcgaatgattagaggccttggggttgaaacaaccttaac
ctattctcaaactttaaatatgtaagaagtccttgttacttaattgaacgtggacattcg
aatgtaccaacactagtgggccatttttggtaagcagaactggcgatgcgggatgaaccg
aacgtgaagttaaggtgccggaatct----aggctcatcagacaccacaaaaggtgttag
ttcatctagacagcaggacggtggccatggaagtcggaatccgctaaggagtgtgtaaca
actcacctgccgaatgaactagccctgaaaatggatggcgcttaagcctagtacccatac
ttcaccgccagggtagaaacgatgccctggcgagtaggcaggcgtggaggtcagtga
>Lachnum_palmae_TNS_F__24600
----------------------------------------gcctcagtaacggcgagtg
aagcggcaacagctcaaatttgaaatctggctccttcggggcccgagttgtaatttgtag
aagatgctttgagggcggc-gcggcctaagttccttggaacaggacgtcagagagggtga
gaatcccgtctggtcaacg----cctaactcgtgtaaagctctttcgacgagtcgagttg
tttgggaatgcagctcaaaatgggtggtaaatttcatctaaagctaaatactggccagag
accgatagcgcacaagtagagtgatcgaaagatgaaaagcactttggaaagagagttaaa
cagtacgtgaaattgttgaaagggaagcgttggcacccagacttgt-cccgaacgctcag
cgggggttcgcccccgtgtattcgtttgtg-gcaggccagcatcggttctggtggcggga
taaa-cccaggggaacgtggctcttc----ggagtgttatagcccttg--gccataccgc
ctaccgggaccgaggaccgcgctttggctaggatgctggcgtaatgggtgtcaacggccc
gtcttgaaacacggaccaaggagtctaacatctatgcaagtatttgggtgctaaaaccca
tatgcgcaatgaaagtgaacggaggtaggcgccctt-aagggtccactatcgaccgatcc
tgatgtcttcggatggatttgagtatgagcatagctgttgggacccg-aaagatggtgaa
ctatgcctaaatagggtgaagccagaggaaactctggtggaggctcgcagcggttctgac
gtgcaaatcgatcgtcaaatttgggcataggggcgaaagactaatcgaactat-------
------------------------------------------------------------
------------------------------------------------------------

------------------------------------------------------------

------------------------------------------------------------

------------------------------------tttgctactaggtagttaagagaa

ttataatctagtccctcttgcgtggggcgacactgtcaaattgcggggacgtcctgttat

gctaaactactgcaacattgtggaaacacgtgtgcgcaccagggt---aatgacctgggg

atagtaacaacgtttagagtagggataattcgcagccaagccctaaa----ggtttttg

aacccatgggtgcagttcacagactaaatgtcagtgggcctcgaggcacgctcgaggctt

aagttatagtcggaccgtcggtaaaccgaagagcaagttgggatcttgatagcatgcaag

ctataatattcacgtattatatggctgt-caagagaaatcttgctaaaaactttgtttta

att-------gcgtcgaagcagtccgaaaa------------------------------

------------------------------------------------------caatag

>Lachnum_rachidicola_NBRC_114473

------------------------------------------------------------

------------------------------------------------------------

------------------------------------------------------------

------------------------------------------------------------

------------------------------------------------------------

------------------------------------------------------------

--------tgaaactcccgaaggacaagcttgtgggcttgtcaagaacttggctcttatg

tgttatgtcactgttggtacgcctagtgatcctatcatcgagttcatgatacaaagaaat

atggaagtcctggaagaatatgagcccttgcggtcacctaatgccaccaaagtcttcgtt

aatggcgtttgggttggtgtgcatcgggaccctgcacatcttgtgaggacagtgcagcat

ctgagacgatcacatttgatatctcacgaagtttctttgattcgggatattcgagaccga

gaattcaagatcttcacagacgcaggcagagtgtgcagacctctcttcgtcattgacaac

gacgttgacagtgccaacaagggcaatttggttttgagcaaggagcatatccgacggctt

gaagaagaccaaaccatgcccgcgaatatggactcggaacagaaagccaatgcaggctac

tacggattccaaggcttaattaatgatggtgtggtcgaatatgtggatgctgaagaagaa

gagacagtcatgatcgtgatgacaccggaggacttggatatttctcgtcaactgcaagcg

ggctaccaaatcagaccagatgagagcggagatctgaacaagcgtgtgaaagcgcccatg

aatccaaccgcgcacatatggacccattgcgagattcatccgagtatgattttgggaatt

tgcgcaagcatcattcccttcccggatcacaaccaggtaagacctggagccgatttttgt

tgatcatttcgctgatnattagacagtctcctcgnaacacctacca--------------

-----

>Lachnum_virgineum_TNS__F__16583

------------------------------------------------------------

------------------------------------------------------------

------------------------------------------------------------

------------------------------------------------------------

------------------------------------------------------------

------------------------------------------------------------

-------------------------------gtgggctcgtc-agaacctggctcttatg

tgctacgtcacagtcggtacgcctagcgaccctattattgagttcatgatccaaagaaac

atggaagtcctggaggaatacgagcctttgcggtcgccgaatgcaacgaaagtcttcgtg

aatggtgtatgggttggtgtgcatcgggacccggcacatctcgtgaaaacagtgcagcat

ctgagacgatcacatttaatctcccacgaagtttccctcattcgggatattcgagaccga

gagtttaagatcttcacggacgcaggccgtgtctgtagacctcttttcgtcgttgataac

gatgttgatagtgcgaacaagggcaatttggttttgaacaaggaccacatccgaaaactt

gaagaagatcagacgatgcccgccaatatggacgccgatcagagaactgatgctggatat

tacggtttccagggtttaattaatgatggtgtggtcgagtacgttgatgccgaagaggag

gagactgttatgatcgtaatgactcctgaggacctggatatttcccgtcaattacaagct

ggtttccagatcagaccggatgaaagcggggatctaaacaaacgtgtgaaagcaccgatg

aaccctactgcacatatctggactcattgcgagattcatccaagtatgattttgggaatt

tgcgcaagcattattcccttcccagatcacaatcaggtaaggactagaactattttactc

ggattcctttgctaa---------------------------------------------

-----

>Lachnum_controversum_MFLU_18__1820

--------------------------------------------------tttgttccgc

agattgacgcaggatgtttacaagtacctgacgaaatgtgtttccgagaacaaggaattc

aacctgacacttggagtgaaatttacaaccctcccaaacggtctcaagtactctttggct

actggaaactggggtgatcaaaaaaaagcagctagctcaaccgctggtgtctctcaggtg

ctgaacagatacacgtttgcctccacgctttcgcatttacgccgtacaaatacacctatt

gggcgtgatggaaagatcgccaagccgcgtcaattgcacaatactcactggggtcttgtc

tgtcctgccgaaactcctgaaggacaagcatgtggacttgtcaagaatttggctctcatg

tgttatgtcacagttggtacgcccagcgaccctatcatcgaattcatgatccaaagaaac atggaagttctagaagaatatgagcctttgcggtctccgaatgcaaccaaagtcttcgtc

aatggtgtgtgggttggtgtgcatcgggaccctgcacacctcgtaaaaacggtgcagcat

ctgcgacgatcacatttgatctctcacgaagtgtctttgattcgtgatattcgagacaga

gaattcaagattttcaccgatgcaggtcgtgtttgcagacctctcttcgttattgacaac

gacgttgacagtactaataagggcaatttggttctgagcaaggagcacatccgacggctt

gaagaggaccagacaatgccggctaatatggactccgaacagaaagctaatgccggctac

tacggtttccaaggcttgattaacgatggtgtagtggaatatgtcgatgccgaagaagag

gagacggtcatgattgtaatgactcctgaggacttggatatctctcgtcaattgcaagct

ggttaccagatcagaccggatgaaagcggggatttgaacaagcgtgtgaaagcaccgatg

aatccaactgcacatatctggacccattgtgagattcatcccagtatgatcttgggaatt

tgcgcaagcattatccccttcccagatcataaccaggtaaggactataattggattccgt

tgaaacttctgc------------------------------------------------

-----

>DSM10201

------------------------------------------------------------

-----------------------gtacctaacgaaatgtgttcaagagaataaggaattc

aacctcacccttggtgtgaaatctacaactctcacaaacggtctcaagtactctctggca

actggaaactggggtgaccagaagaaagcagctagctcgaccgctggtgtgtctcaggtg

ctgaacagatatacatttgcctccacactctcgcatttacgccgtacgaatacgcctatt

gggcgagatggaaagatcgccaaaccgcgtcaactgcacaacactcattggggtctcgtt

tgtcccgccgaaactcccgaaggacaggcttgtgggcttgtcaagaacttggctctcatg

tgttatgtcaccgtgggtactcctagtgaccctattattgagttcatgatccaacgaaac

atggaagttttggaggaatatgagcctttgcggtctccgaatgcgaccaaagtcttcgtc

aatggtgtatgggttggtgtgcatcgggaccctgcacatctcgtaaaaacagtgcagcat

ctgagacgatcacatttgatttctcacgaagtctctttaattcgagatattcgagaccga

gaattcaaaatcttcacagacgcaggccgcgtctgtagacctctcttcgtcattgacaac

gatgttgatagtacaaacaagggaaatttggttttgaacaaggaccacatccgacgactt

gaagatgatcagacgatgcccgcaaatatggactcggaacagaaagccaacgctggttat

tatggtttccagggtttaattaacgatggtgtggtcgagtatgttgatgctgaagaagag

gagactgtaatgatcgtgatgactcctgaggacctggacatttcccgtcaattacaagct

ggttaccagatcagaccggatgaaagtggagatctgaacaagcgtgtgaaagcaccgatg

aatccaactgcacacatctggacccattgcgagatccatccaagtatgattttgggaatt tgcgcaagcattattcccttcccagatcacaaccaggtaagggctagaattaact-----

------------------------------------------------------------

-----

>ATCC36554

------------------------------------------------------------

------------------------------------------------aataaggaattc

aaccttacccttggtgtgaaatctacaactctcacaaacggtctcaagtactctctggca

actggaaactggggtgaccagaagaaagcagctagctcgaccgctggtgtctctcaggtg

ctgaacaggtatacctttgcctccacactctcgcatttacgccgtacgaatacacctatt

gggcgagatggaaagattgccaagccgcgccaattgcacaacactcattggggtcttgtc

tgtcctgccgaaactcctgaaggacaagcttgtgggcttgtcaagaacttggctctcatg

tgttacgttacagttggtacgcctagtgaccctattattgagttcatgatccaaagaaac

atggaagttctggaggaatatgagcctttgcggtctccgaatgcgaccaaagtcttcgtc

aatggtgtatgggttggtgtacatcgggaccctgcacatctcgtaaaaacagtgcagcat

ctgagacgatcgcatttaatctcccacgaagtttccctgattcgggatattcgagaccga

gagtttaagatcttcacagatgcaggccgtgtctgtagacctcttttcgtcattgacaac

gatgttgatagtgcacagaagggcaatttggttctaaacaaggagcacatccgaaaactt

gaagatgatcagacaatgcccgcaaatatggactcggagcagaaagccaacgctggttat

tatggtttccaaggtttaatcaacgatggtgtggtcgagtacgtcgatgctgaagaagag

gagactgtaatgatcgtgatgactcctgaagacttggatatttcccgtcaattgcaggct

ggctaccagatcagaccggatgaaagtggggacttgaacaagcgtgtgaaagcaccgatg

aatccaactgcacacatctggacccattgcgagattcatccaagtatgattttgggaatt

tgcgcaagcattattcccttcccaga----------------------------------

------------------------------------------------------------

-----

>Lachnum_soppittii_FC__2160

------------------------------------------------------------

------------------------------------------------------------

------------------------------------------------------------

------------------------------------------------------------

------------------------------------------------------------

------------------------------------------------------------

-----------------------------------------------tttggctctcatg

tgttacgtcacagttggtacgcccagcgaccctattattgagttcatgatccaaagaaac

atggaagttttggaggaatatgagcctttgcggtctccaaatgcgaccaaagtcttcgtc

aatggtgtatgggttggtgtgcatcgggaccccgcacatctagtgaaaacagtgcagcat

ctgagacgatcgcacttgatctcccacgaagtctccttaattcgggatatccgagaccga

gaattcaagatcttcacagacgcaggtcgtgtctgtagaccactcttcgtcattgacaac

gatgttgagagtaataacaagggcaatttggtcttgaccaaggaccatattcgaagactt

gaagacgatcagacgatgcccgcaaatatggatgcggagcagagagaggcagctaactat

tttggtttccaaggtttaatcaatgagggtgtggtcgagtatgtagatgctgaagaagag

gagactgtcatgatcgtgatgactccagaggacctggacatttcccgccaattgcaagct

gggttgcaaatcagaccagatgaaagcggagacttgaacagacgtgtgaaagcaccgatg

aatccaacagcacacatctggacccattgtgagatccatccaagtatgattttgggaatt

tgcgcaagcattatcccctttccagatcacaaccaggtacggagcagtattgattttggt

tgattcttttgctaatggacatcc------------------------------------

-----

>Lachnum_palmae_TNS_F__24600

------------------------------------------------------------

------------------------------------------------------------

------------------------------------------------------------

------------------------------------------------------------

------------------------------------------------------------

------------------------------------------------------------

tgtcctgctgaaactcctgaaggacaagcttgtggactagtcaagaacctggcactgatg

tgctatgtgacagtcggaacaccgagtgaacccataattgagttcatgatccaaagaaat

atggaagtcttagaagagtacgagcctttacggtctccaaacgcaaccaaagtctttgtt

aatggtgtctgggttggcgtgcaccgcgatcctgctcatcttgtgcgaacagtgcagcat

ctgagacgatcacatctgatctctcatgaagtctctttgatcagagatattcgagataga

gagttcaagattttcaccgatgcaggccgtgtctgtagaccactgttcgtcattgacaac

gatattgacagtgcgaacaaaggtaatttggttttgaataaggagcacatccggcggctt

gaagacgaccaggcgatgcctgccaatatggacgctgatcagagagccagtgctggttac

tatggattccagggcttgatcaatgatggtgttgttgagtacgtggatgctgaggaagaa

gagactgttatgattgtcatgacacctgaagacttggatatttctcgtcagctccaagcc gggtatcagattagaccagatgagagcggggatctcaataagcgtgtgaaagccccaatg

aacccaactgcacacatctggacgcactgcgagattcatccaagtatgattctagggatt

tgcgcaagcatcatccccttccccgatcacaaccaggtaagaatcaagctaagctttgag

gacgttgctaatgactttagtccc------ctcgtaatacttaccagtctgctatgggta

agcaa

>Lachnum_abnorme_KUS__F52080

------------------------------------------------------------

------------------------------------------------------------

------------------------------------------------------------

------------------------------------------------------------

------------------------------------------------------------

------------------------------------------------------------

------------actcccgagggccaagcttgtggactggtcaagaatttggctcttatg

tgctatgtcacagttggtacgcccagcgacccgattattgagttcatgattcaaaggaac

atggaagtattggaagaatatgaacctctgcggtccccaaatgctacaaaggtctttgtc

aacggcgtttgggttggtgtgcatcgcgatcccgctcatctcgtgaggacagtgcagcat

ctgagacgatcgcatctaatctctcacgaagtttctttgatccgagacattcgtgataga

gaattcaagatctttactgatgcaggccgcgtctgccgaccgctcttcgtcgttgataat

gatgtcgacagtcctaacaaaggcaatttggttttgaataaggagcacatccgacggctt

gaagatgatcaaactatgcctgccaatatggacttagaacagagggctagtgcaggctac

tatggtttccagggtctaatcaatgatggtgtggtcgagtatgtggacgccgaggaagag

gagactgttatgattgtcatgactcccgaggacttggatatttctcgacagttgcaagct

ggttatcagatcagaccagatgaaagcggggatctgaataagcgtgtgaaagcaccgatg

aacccaactgcacatatctggacgcattgcgagattcatccaagtatgatattggggatt

tgcgcaagcattatccccttcccggatcacaaccaggtaataaatccagaacttattcag

agtcccctctctaattgtatc---------------------------------------

-----

>Proliferodiscus_sp._KUS__F52660

aagaagcgtttggatcttgctggacctctccttgctaagctgtttcgaagtctcttccgt

aggctgacaacggatgtctacaagtacctcacgaaatgtgtctctgagaacaaagagttc

aatctcacgctcggagtgaagtccactactctcacgaatggtcttaaatattctttggcc

accgggaactggggtgaccagaagaaagcggcaagctccacggctggtgtctctcaggtg ctgaacagatatacctttgcttccacactttcgcatttgcgtcgcacaaatacaccaatt

gggcgtgatggaaagattgctaagccgcgacagctgcataacactcactggggtctggtg

tgtcctgctgagactcctgaagggcaagcttgtgggcttgtcaagaatttggctcttatg

tgctacgtcacagtcggtacgcctagcgagcctatcattgagttcatgattcaaagaaat

atggaagttttggaggaatacgagcctctgcggtctcccaatgcaacgaaggtcttcgtc

aacggtgtctgggttggtgtgcaccgcgatcctgcacatctggtgaggacggtgcagcat

ttgaggcgatcgcacttgatctctcacgaagtctctctgattcgagatattcgagacaga

gaattcaagatcttcactgatgcaggccgagtctgtagaccactgttcgtcattgacaat

gatgttgacagtccgaacaaaggcaatctggtcttgaataaggaccacattcgacggctt

gaggatgaccagacgatgcctgccaatatggactcagatcagaaggctaatctaggttat

tatggtttccagggcttgatcaatgatggtgttgttgagtacgttgatgctgaggaagag

gagacagtcatgatcgtgatgactcctgaggacttggacatttctcgccaactgcaagcc

gggtatcagataaggccagatgagagcggcgatctgaacaagcgtgtgaaagcgccaatg

aacccgactgcacacatctggacgcattgcgagattcatcctag----------------

------------------------------------------------------------

------------------------------------------------------------

-----

>Proliferodiscus_sp._TNS__F17436

------------------------------------------------------------

------------------------------------------------------------

------------------------------------------------------------

------------------------------------------------------------

------------------------------------------------------------

------------------------------------------------------------

----------agactcctgaagggcaagcttttgggcttgtcaagaatttggctcttatg

tgctacgtcacagtcggtacgcctagcgagcctatcattgagttcatgattcaaagaaat

atggaagttttggaggaatacgagcctctgcggtctcccaatgcaacgaaggtcttcgtc

aacggtgtctgggttggtgtgcaccgcgatcctgcacatctggtgaggacggtgcagcat

ttgaggcgatcgcacttgatctctcacgaagtctctctgattcgagatattcgagacaga

gaattcaagatcttcactgatgcaggccgagtctgtagaccactgttcgtcattgacaat

gatgttgacagtccgaacaaaggcaatctggtcttgaataaggaccacattcgacggctt

gaggatgaccagacgatgcctgccaatatggactcagatcagaaggctaatctaggttat

tatggtttccagggcttgatcaatgatggtgttgttgagtacgttgatgctgaggaagag
gagacagtcatgatcgtgatgactcctgaggacttggacatttctcgccaactgcaagcc
gggtatcagataaggccagatgagagcggcgatctgaacaagcgtgtgaaagcgccaatg
aacccgactgcacacatctggacgcattgcgagattcatcctagtatgattttggggatt
tgcgcaagcattatccccttcccagatcacaatcaggtaagaagatttgtccaatttgcg
tgctcctttactgat---------------------------------------------
-----
>Lachnum_fuscescens_FC__2200
------------------------------------------------------------
------------------------------------------------------------
------------------------------------------------------------
------------------------------------------------------------
------------------------------------------------------------
------------------------------------------------------------
-----------------------------------gcttgtcaagaacctggcgctcatg
tgctacgtcacagtcggtacacccagcgagcctattattgagttcatgatccaaagaaac
atggaggttttggaggagtacgagccgttgaggtccccgaatgccacgaaggtgtttgta
aatggtgtttgggtgggtgtgcacagagaccctgcgcatttggtgaagacagtgcaacat
ctgagacgatcgcatttgatctcccacgaagtctcccttattcgagacatcagagaccga
gaattcaagattttcacagacgcaggacgagtctgcagaccgcttttcgtcattgataac
gatgttgacagtgccaacaagggcaatttggtcttgaacaaggagcacattaggcgactt
gaagaagaccaaactatgccagccaacatggacgccgaacagaaggaaaactccggctac
ttcggattccagggcttgatcgatggtggagtggtcgagtatgtagatgctgaagaagaa
gaaaccgtgatgattgtgatgaccccagaagatttggatatctctcgtcaacttcaggct
ggttaccaaatcagacctgacgatagtggggacttgaacaagcgtgtgaaagcaccgatg
aatcctacagcacatatttggacacattgcgagattcatccaagtatgattttgggaatt
tgcgcaagcattatccctttcccagatcacaatcaagtaaggatctcacagagttt--ga
tgttctttatactaatgtgcc---------------------------------------
-----
>Brunnipila_fuscescens_KUS__F52031
---------ctagatcttgcaggacctcttcttgctaaacttttccgaaatctcttccgc
agattaacaggagatgtgttcaagtatttgcagaaatgtgtggcggataacaaggagttc

aacttgactctgggtgtcaaatccaccaccctcacaaatggtctcaaatattctttggct
accggcaactggggcgaccaaaagaaggcggccagctcaaccgctggtgtgtcccaagtg
ctaaacagatacactttcgcttctacactatcccatttacggcgtacaaacacccccatt
ggccgtgatggaaaaattgctaaaccgcgacaacttcacaacactcattggggtcttgtc
tgtcctgccgaaacgcctgaaggacaagcttgtgggcttgtgaagaatttggcgctcatg
tgctacgtcacagtcggtacacccagtgaacctatcattgagttcatgatccaaagaaac
atggaggtcttagaggaatacgagccgttgcggtcgccgaatgccacaaaggttttcgtg
aacggtgtttgggttggtgttcatcgagatccagcgcatctggtgaggacagtgcagcat
ttgagacgatcacatttgatttctcacgaagtttccctgatccgagacattagagatcga
gagttcaagatttttacagatgcaggacgagtctgcagaccgctcttcgtcattgacaat
gatgttgacagtgctaacaagggcaatctggtcttgaacaaggagcacattaggcgactt
gaagaagaccagactatgccagccaacatggaccccgaacaaaaggccaactccggctac
ttcggtttccaaggcttgatcaatgacggtgttgttgaatatgtggacgctgaagaagag
gagactgttatgattgtgatgacccctgaagatttggatatctctcgtcaactacaggcc
gggtaccaaattcgaccagacgatagcggggacctgaacaagcgtgtaaaagcaccaatg
aaccctacagcacacatctggacgcattgcgagatccatcctagtatgattctgggaatt
tgcgcgagcattattcctttcccagatcacaaccaagtaaggagtgaacagagcttt--ga
tatatgttttgcta----------------------------------------------
-----
>Trichopeziza_sulphurea_KUS__F52218
------------------------------------------------------------
------------------------------------------------------------
------------------------------------------------------------
------------------------------------------------------------
--------------------------------------gcgaagaacaaacactcctatt
ggtcgagatggaaagattgccaaacctcgtcaattgcacaacactcattggggtctggta
tgtcctgccgagactcctgaggggcaagcttgtggtctggtcaagaatctggcactcatg
tgctatgtcactgttggtacacctagtgacccaatcattgagtttatgatccagagaaac
atggaagttctggaggaatacgaacctttacgttctccaaatgccaccaaggtttttgtg
aatggtgtttgggttggtgtacacagagaccctgcccatcttgtcaggacggtgcaacat
cttcgtcgttctcacttgatctctcacgaagtctccttaattcgagatattcgagatcga
gaattcaaaatcttcacagacgcgggtagagtgtgtagacccttgttcgttattgacaac

gatcttgatagcccaaacaagggaaatctggtactcaataaaatgcacattggacgatta
gaagacgatcaaacaatgcctgcaaatatggacatggaacaaagagtcactcagggtcat
tttggtttccaaggtctcatcaatgaaggtgtggttgaatatgttgatgcggaggaagaa
gagacggtaatgattgtgatgacccccgaagatttggatatctcgcgtcaactccaggca
ggttatcaaattagacctgatgagagtggggatttgaacaaacgtgtgaaagcacctatg
aatccaacagctcatatctggactcattgcgagattcatccaagtatgattctgggaatt
tgcgcaagcattattcctttcccagatcacaatcaggtaagatcattcatgctaagtttc
tg----------------------------------------------------------
-----
>Arachnopeziza_aurelia_TNS__F11211
------------------------------------------------------------
------------------------------------------------------------
------------------------------------------------------------
------------------------------------------------------------
------------------------------------------------------------
------------------------------------------------------------
--------------------------------------------------agctctcatg
tgctacgttacggttggaacgcctagcgatcccattattgagtttatgattcagcgtaac
atggaagttcttgaagaatacgagccactgagatccccaaatgctacaaaagttttcgtc
aacggtgtctgggttggggtccacagagatcctgcccatctggttcagacagtgcaaaat
cttcgaagatctcacttgatctctcacgaggtttcattgattcgagatattcgagaccga
gaattcaagattttcacagatgctggcagagtgtgtcggccacttttcgtcattgaaaat
ggtattgacaatcctaagaaggggcaactagttcttaataaggatcatattcgcagactt
gagcttgatcagacaatg---agtggaatggaccaagatacccgattggctaatggatat
ttcggttttcagggtctaatcaactcgggcgtggttgaatatttggatgccgaagaagaa
gagactgttatgattgtcatgacccctgaggacttagatatctcacgacaactgcaagct
ggatacactcttgaacccgacaccagtggagatatgaataaaagagtgaaggctcctatg
aatcctacagcgcatatgtggacacattgtgagattcatcctagtatgattttgggaatt
tgtgcaagcattattcccttcccagatcataatcaagtaagtacactgt-----------
------------------------------------------------------------
-----
>Arachnopeziza_aurata_TNS__F11212

------------------------------------------------------------

------------------------------------------------------------

------------------------------------------------------------

------------------------------------------------------------

------------------------------------------------------------

------------------------------------------------------------

------------------------------------------------------------

------------------------------------------------------------

------------------------------------------------------------

------------------------------------------------------------

----gaagatctcacttgatatctcacgaagtctctttaatcagagatattcgagaccga

gaatttaaaatcttcacggatgcaggcagagtatgccgacctctttttgttattgaaaat

ggtcatgacaatcccaacaagggaaacttggttttgaacaaagatcatattcgtaagctc

gagttagatcagaccttg---gcaggaatggatcaagaaactcgattggcgaatggatac

tttggcttccaaggtctaatcaactctggagttgtcgaatatctggatgctgaggaagag

gaaacggtcatgatagtcatgactcctgaggatttggatatatcacggcaattacaagct

ggcttcaagattcaacctgacgatagtggggatatgaataagagagttaaggctcctatg

aaccctactgcacatatgtggacgcattgtgagattcatccaagtatgatcttggggatt

tgcgcaagcattatccccttcccagatcataatcaagtaagtccattgttggtcat--ga

tgtctaatgcatgcta--------------------------------------------

### Alignment of the LSU sequences used in the phylogenetic study

>Lachnum_rachidicola_NBRC_114473

tagcatatcaataagcggaggaaaagaaaccaacagggattacctcagtaacggcgagtg

aagcggtaacagctcaaatttgaaatctggctctttcagggtccgagttgtaatttgtag

aagatgctttgggtgtggtccgggtctaagttccttggaacaggacgtcatagagggtga

gaatcccgtatgtgactcggtaccttcgcccgtgtaaagctctttcgacgagtcgagttg

tttgggaatgcagctcaaaatgggtggtaaatttcatctaaagctaaatattggccagag

accgatagcgcacaagtagagtgatcgaaagatgaaaagcactttggaaagagagttaaa

cagtacgtgaaattgttgaaagggaagcgcttgcaaccagacttagcggcggctgatcat

ccggggttctccccggtgcactcggtcgtctctaggccagcatcggtttgggtggtggga

taaaggccttggtaatgtagcttctctcggggagtgttatagacctcggtgcaatgccgc

ctacccggaccgaggaccgcgcttcggctaggatgctggcgtaatggttgtaagcgaccc gtcttgaaacacggaccaaggagtctaacatctatgcgagtatttgggtgtc-aaaccca

tatgcgtaatgaaagtgaacggaggtgagaaccctt-aagggtgcatcatcgaccgatcc

tgatgtcttcggatggatttgagtaagagcatagctgttgggacccg-aaagatggtgaa

ctatgcctaaatagggtgaagccagaggaaactctggtggaggctcgcagcggttctgac

gtgcaaatcgatcgtcaaatttgggtatagggcgaaagactaatcgaaccat-------

------------------------------------------------------------

------------------------------------------------------------

------------------------------------------------------------

------------------------------------------------------------

------------------------------------ctagtagctgg-------------

------------------------------------------------------------

------------------------------------------------------------

------------------------------------------------------------

------------------------------------------------------------

------------------------------------------------------------

------------------------------------------------------------

------------------------------------------------------------

---------------------------------------------------------

>Lachnum_virgineum_TNS__F__16583

---------------------------------------------cagtaacggcgagtg

aagcggtaacagctcaaatttgaaatctggctctttcagggtccgagttgtaatttgtag

aagatgctttgggtgtggccctggtctaagttccttggaacaggacgtcatagagggtga

gaatcccgtatgtgactaggtgctttcgcccgtgtaaagctctttcgacgagtcgagttg

tttgggaatgcagctcaaaatgggtggtaaatttcatctaaagctaaatattggccagag

accgatagcgcacaagtagagtgatcgaaagatgaaaagcactttggaaagagagttaaa

cagtacgtgaaattgttgaaagggaagcgcttgcaaccagacttagttgcggccgatcat

ctagggttctccctggtgcactcggtcgtatctaggccagcatcggtttgggtggtggga

taaaggccttgggaatgtagcttctttcggggagtgttatagccctcggtgcaatgccgc

ctacctggaccgaggaccgcgcttcggct-------------------------------

------------------------------------------------------------

------------------------------------------------------------

------------------------------------------------------------

------------------------------------------------------------

------------------------------------------------------------

------------------------------------------------------------

------------------------------------------------------------

------------------------------------------------------------

------------------------------------------------------------

------------------------------------------------------------

------------------------------------------------------------

------------------------------------------------------------

------------------------------------------------------------

------------------------------------------------------------

------------------------------------------------------------

------------------------------------------------------------

------------------------------------------------------------

---------------------------------------------------------

>DSM10201

---------------------------------------------------cggcgagtg

aagcggtaacagctcaaatttgaaatctggctcttttagggtccgagttgtaatttgtag

aagatgctttgggtgtggccctggtctaagttccttggaacaggacgtcatagagggtga

gaatcccgtatgtgactaggtgtcttcgcccgtgtaaagctctttcgacgagtcgagttg

tttgggaatgcagctcaaaatgggtggtaaatttcatctaaagctaaatattggccagag

accgatagcgcacaagtagagtgatcgaaagatgaaaagcactttggaaagagagttaaa

cagtacgtgaaattgttgaaagggaagcgcttgcaaccagacttagttgcggccgatcat

ctagggttctccctggtgcactcggtcgtatctaggccagcatcggtttgggtggtggga

taaaggccttgggaatgtagcttctttcggggagtgttatagccctcggtgcaatgccgc

ctacctggaccgaggaccgcgcttcggctaggatgctggcgtaatggttgtaagcgaccc

gtcttgaaacacggaccaaggagtctaacatctatgcgagtatttgggtgtt-aaaccca

tatgcgtaatgaaagtgaacggaggtgagaaccctt-aagggtgcatcatcgaccgatcc

tgatgtcttcggatggatttgagtaagagcatagctgttgggacccg-aaagatggtgaa

ctatgcctaaatagggtgaagccagaggaaactctggtggaggctcgcagcggttctgac

gtgcaaatcgatcgtcaaatttgggtataggggcgaaagactaatcgaaccat-------

------------------------------------------------------------

------------------------------------------------------------
------------------------------------------------------------
------------------------------------------------------------
------------------------------------ctagtagctggttcctgccgaag-
---------------tttccctcaggatagcagtgttgaatt------------------
----cagttttatgaggtaaagcgaatgattagaggccttggggttgaaacaaccttaac
ctattctcaaactttaaatatgtaagaagtccttgttacttaattgaacgtggacattcg
aatgtaccaacactagtgggccatttttggtaagcagaactggcgatgcgggatgaaccg
aacgtgaagttaaggtgccggaatat----acgctcatcagacaccacaaaaggtgttag
ttcatctagacagcaggacggtggccatggaagtcggaatccgctaaggaatgtgtaaca
actcacctgccgaatgaactagccctgaaaatggatggcgcttaagcgtattacccatac
ttcaccgccagggtagaaacgatgccctggcgagtaggc------------------
>ATCC36554
------------------------------------------------------------
--gcggtaacagctcaaatttgaaatctggctctttcagggtccgagttgtaatttgtag
aagatgctttgggtgtggccctggtctaagttccttggaacaggacgtcatagagggtga
gaatcccgtatgtgactaggtgctttcgcccgtgtaaagctctttcgacgagtcgagttg
tttgggaatgcagctcaaaatgggtggtaaatttcatctaaagctaaatattggccagag
accgatagcgcacaagtagagtgatcgaaagatgaaaagcactttggaaagagagttaaa
cagtacgtgaaattgttgaaagggaagcgcttgcaaccagacttagttgcggccgatcat
ctagggttctccctggtgcactcggtcgtatctaggccagcatcggtttgggtggtggga
taaaggccttgggaatgtagcttctttcggggagtgttatagccctcggtgcaatgccgc
ctacctggaccgaggaccgcgcttcggctaggatgctggcgtaatggttgtaagcgaccc
gtcttgaaacacggaccaaggagtctaacatctatgcgagtatttgggtgtt-aaaccca
tatgcgtaatgaaagtgaacggaggtgagaaccctt-aagggtgcatcatcgaccgatcc
tgatgtcttcggatggatttgagtaagagcatagctgttgggacccg-aaagatggtgaa
ctatgcctaaatagggtgaagccagaggaaactctggtggaggctcgcagcggttctgac
gtgcaaatcgatcgtcaaatttgggtataggggcgaaagactaatcgaaccat-------
------------------------------------------------------------
------------------------------------------------------------
------------------------------------------------------------
------------------------------------------------------------

------------------------------------ctagtagctggttcctgccgaag-

---------------tttccctcaggatagcagtgttgaatt------------------

----cagttttatgaggtaaagcgaatgattagaggccttggggttgaaacaaccttaac

ctattctcaaactttaaatatgtaagaagtccttgttacttaattgaacgtggacattcg

aatgtaccaacactagtgggccatttttggtaagcagaactggcgatgcgggatgaaccg

aacgtgaagttaaggtgccggaatat----acgctcatcagacaccacaaaaggtgttag

ttcatctagacagcaggacggtggccatggaagtcggaatccgctaaggaatgtgtaaca

actcacctgccgaatgaactagccctgaaaatggatggcgcttaagcgtattacccatac

ttcaccgccagggtagaaacgatgccctggcgagtaggcaggcgtgggggtca----

>Lachnum_soppittii_FC__2160

--------------------------------------attacctcagtaacggcgagtg

aagcggtaacagctcaaatttgaaatctggctctttcagggtccgagttgtaatttgtag

aagatgctttgggtgtggcccgggtctaagttccttggaacaggacgtcatagagggtga

gaatcccgtatgtgacccggcgctttcgcccgtgtaaagctctttcgacgagtcgagttg

tttgggaatgcagctcaaaatgggtggtaaatttcatctaaagctaaatattggccagag

accgatagcgcacaagtagagtgatcgaaagatgaaaagcactttggaaagagagttaaa

cagtacgtgaaattgttgaaagggaagcgcttgcaaccagacttaggcgcggccgatcat

ccagggttctccctggtgcactcggtcgtctctaggccagcatcggtttgggtggtggga

taaaggccttgggaatgtagcttctttcggggagtgttatagccctcggtgcaatgccgc

ctacctggaccgaggaccgcgcttcggctaggatg-------------------------

------------------------------------------------------------

------------------------------------------------------------

------------------------------------------------------------

------------------------------------------------------------

------------------------------------------------------------

------------------------------------------------------------

------------------------------------------------------------

------------------------------------------------------------

------------------------------------------------------------

------------------------------------------------------------

------------------------------------------------------------

------------------------------------------------------------

------------------------------------------------------------

------------------------------------------------------------

------------------------------------------------------------

------------------------------------------------------------

------------------------------------------------------------

---------------------------------------------------------

>Lachnum_controversum_MFLU_18__1820

-----------aatagcggagaaaagaaaccaacagggattacctcagtaacggcgagtg

aagcggtaacagctcaaatttgaaatctggctctttcagggtccgagttgtaatttgtag

aagatgctttgggtgtggcccgggtctaagttccttggaacaggacgtcatagagggtga

gaatcccgtatgtgactcggcgccttcgcccgtgtaaagctctttcgacgagtcgagttg

tttgggaatgcagctcaaaatgggtggtaaatttcatctaaagctaaatattggccggag

accgatagcgcacaagtagagtgatcgaaagatgaaaagcactttggaaagagagttaaa

cagtacgtgaaattgttgaaagggaagcgcttgcaaccagacttagttgctgtcgatcat

ctcgggttctcccgggtgcactcggcagtgtctaggccagcatcggtttgggtggtggga

taaaggccttgggaatgtagcttctttcggggagtgttatagccctcggtgcaatgccgc

ctacccggaccgaggaccgcgcttcggctaggatgctggcgtaatggttgtaagcgaccc

gtcttgaaacacggaccaaggagtctaacatctatgcgagtatttgggtgtc-aaaccca

tatgcgtaatgaaagtgaacggaggtgagaaccctt-aagggtgcatcatcgaccgatcc

tgatgtcttcggatggatttgagtaagagcatagctgttgggacccg-aaagatggtgaa

ctatgcctaaatagggtgaagccagaggaaactctggtggaggctcgcagcggttctgac

gtgcaaatcgatcgtcaaatttgggtataggggcgaaagactaatcgaac-at-------

------------------------------------------------------------

------------------------------------------------------------

------------------------------------------------------------

------------------------------------------------------------

------------------------------------ctag-agctggtcc----------

------------------------------------------------------------

------------------------------------------------------------

------------------------------------------------------------

------------------------------------------------------------

------------------------------------------------------------

------------------------------------------------------------

------------------------------------------------------------

---------------------------------------------------------

>Trichopeziza_sulphurea_KUS__F52218

------------------------------------------------------------

--------------caaatttgaaatctggccctttcagggtccgagttgtaatttgtag

aagatgctttgggtgtggctccggtctaagttccttggaacaggacgtcatagagggtga

gaatcccgtatgtgaccggtggctttcgcccgtgtaaagctctttcgacgagtcgagttg

tttgggaatgcagctcaaaatgggtggtatatttcatctaaagctaaatattggccagag

accgatagcgcacaagtagagtgatcgaaagatgaaaagcactttggaaagagagttaaa

cagtacgtgaaattgttgaaagggaagcgcttgcaaccagactcgcgcgccgttgatcat

ccggggttctccccggtgcactcggcggcgctcgggccagcatcagtttgggtggtggga

taaaggccttgggaatgtagctcccctcggggagtgttatagccctcggtgcaatgccac

ctacccggactgaggaccgcgcttcggctaggatgctggcgtaatggttgtaagcgaccc

gtcttgaaacacggaccaaggagtctaacatctatgcgagtgtttgggtgtc-aaaccca

tacgcgtaatgaaagtgaacggaggtgagaacccctt-aagggtgcatcatcgaccgatcc

tgatgtcttcggatggatttgagtaagagcatagctgttgggacccg-aaagatggtgaa

ctatgcctaaatagggtgaagccagaggaaactctggtggaggctcgcagcggttctgac

gtgcaaatcgatcgtcaaatttgggtataggggcgaaagactaatcgaaccat-------

------------------------------------------------------------

------------------------------------------------------------

------------------------------------------------------------

------------------------------------------------------------

------------------------------------ctagtagctggttcctgccgaag-

---------------tttccctcaggatagcagtgttgaatt------------------

----cagttttatgaggtaaagcgaatgattagaggccttggggatgcaacatccttaac

ctattctcaaactttaaatatgtaagaagtccttgttacttaattgaacgtggacattcg

aatgtaccaacactagtgggccatttttggtaagcagaactggcgatgcgggatgaaccg

aacgtgaagttaaggtgccggaatat----acgctcatcagacaccacaaaaggtgttag

ttcatctagacagcaggacggtggccatggaagtcggaatccgctaaggagtgtgtaaca

actcacctgccgaatgaactagccctgaaaatggatggcgcttaagcgtattacccatac

ttcaccgcc------------------------------------------------

>Brunnipila_fuscescens_KUS__F52031

------------------------------------------------taacggcgagtg

aagcggtaacagctcaaatttgaaatctggctctctcagggtccgagttgtaatttgtag

aagatgctttgagtgtggctctagtctaagttccttggaacaggacgtcatagagggtga

gaatcccgtatgtgattaggcgccttcgctcgtgtaaagctctttcgacgagtcgagttg

tttgggaatgcagctcaaaatgggtggtaaatttcatctaaagctaaatattggccagag

accgatagcgcacaagtagagtgatcgaaagatgaaaagcactttggaaagagagttaaa

cagtacgtgaaattgttgaaagggaagcgcttgcaaccagactcgcatgccgtcgatcat

cctgtgttctcactggtgcactcggcggccttcgggccagcatcggtttgggtggtggga

taaaggccttgggaatgtagctcctctcggggagtgttatagccctcggtgcaatgccgc

ctacctggaccgaggaccgcgcttcggctaggatgctggcgtaatggttgtaagcgaccc

gtcttgaaacacggaccaaggagtctaacatctatgcgagtatttgggtgtc-aaaccca

tatgcgtaatgaaagtgaacggaggtgagaaccctt-aagggtgcatcatcgaccgatcc

tgatgtcttcggatggatttgagtaagagcatagctgttgggacccg-aaagatggtgaa

ctatgcctaaatagggtgaagccagaggaaactctggtggaggctcgcagcggttctgac

gtgcaaatcgatcgtcaaatttgggtataggggcgaaagactaatcgaaccat-------

------------------------------------------------------------

------------------------------------------------------------

------------------------------------------------------------

------------------------------------------------------------

------------------------------------ctagtagctggttcctgccgaag-

---------------tttccctcaggatagcagtgttgaatt------------------

----cagttttatgaggtaaagcgaatgattagaggccttggggttgaaacaaccttaac

ctattctcaaactttaaatatgtaagaagtccttgttacttaattgaacgtggacattcg

aatgtaccaacactagtgggccatttttggtaagcagaactggcgatgcgggatgaaccg

aacgtgaagttaaggtgccggaatat----acgctcatcagacaccacaaaaggtgttag

ttcatctagacagcaggacggtggccatggaagtcggaatccgctaaggaatgtgtaaca

actcacctgccgaatgaactagccctgaaaatggatggcgcttaagcgtattacccatac

ttcaccgccagggtagaaacgatgccctggcgagtaggcaggcgtggagg-------

>Trichopezizella_sp._KUS__F52478

------------------------------------------------------------

------------------tttgaaatctggctcttttagggtccgagttgtaatttgtag

aagatgctttgggtgtggctccggtctaagttccttggaacaggacgtcatagagggtga
gaatcccgtatgtgactggttgccttcgcccgtgtaaagctctttcgacgagtcgagttg
tttgggaatgcagctcaaaatgggtggtatatttcatctaaagctaaatattggccagag
accgatagcgcacaagtagagtgatcgaaagatgaaaagcactttggaaagagagttaaa
cagtacgtgaaattgttgaaagggaagcgcttgcaaccagactcgcgcgctgttgatcat
ccggtgttctcaccggtgcactcagcagtgctcgggccagcatcagtttgggtggtggga
taaaggccttgggaatgtagcttctttcggggagtgttatagccctcggtgcaatgccgc
ctacctggactgaggaccgcgcttcggctaggatgctggcgtaatggttgtaagcgaccc
gtcttgaaacacggaccaaggagtctaacatctatgcgagtgtttgggtgtc-aaaccca
tacgcgtaatgaaagtgaacggaggtgagaacccttaaagggtgcatcatcgaccgatcc
tgatgtcttcggatggatttgagtaagagcatagctgttgggacccg-aaagatggtgaa
ctatgcctaaatagggtgaagccagaggaaactctggtgga-gctcgcagcggttctgac
gtgcaaatcgatcgtcaaatttgggtatagggggcgaaagactaatcg-accat-------
------------------------------------------------------------
------------------------------------------------------------
------------------------------------------------------------
------------------------------------------------------------
------------------------------------ctagtagctgt-------------
------------------------------------------------------------
------------------------------------------------------------
------------------------------------------------------------
------------------------------------------------------------
------------------------------------------------------------
------------------------------------------------------------
------------------------------------------------------------
---------------------------------------------------------
>Proliferodiscus_sp._KUS__F52660
------------------------------------------------------------
--------------caaatttgaaatctggctctttcagggtccgagttgtaatttgtag
aagatgctttgggtgtggccccggtctaagttccttggaacaggacgtcatagagggtga
gaatcccgtatgtgactgggtgctttcgcccgtgtaaagctctttcgacgagtcgagttg
tttgggaatgcagctcaaaatgggtggtaaatttcatctaaagctaaatattggccagag

accgatagcgcacaagtagagtgatcgaaagatgaaaagcactttggaaagagagttaaa

cagtacgtgaaattgttgaaagggaagcgcttgcaaccagactcgcatgccgctgatcat

ccggggttctccccggtgcactcggtggtctacgggccagcatcggtttgggtggcggga

taaaggctgtgggaatgtagcttctctcggggagtgttatagcccacggtgcaatgccgc

ctacctggaccgaggaccgcgcttcggctaggatgctggcgtaatggttgtaagcgaccc

gtcttgaaacacggaccaaggagtctaacatctatgcgagtatttgggtgtt-aaaccca

tatgcgtaatgaaagtgaacggaggtaagaaccctt-tagggtgcattatcgaccgatcc

tgatgtcttcggatggatttgagtaagagcatagctgttgggacccgaaaagatggtgaa

ctatgcctaaatagggtgaagccagaggaaactctggtggaggctcgcagcggttctgac

gtgcaaatcgatcgtcaaatttgggtatagggggcgaaagactaatcgaaccat-------

------------------------------------------------------------

------------------------------------------------------------

------------------------------------------------------------

------------------------------------------------------------

------------------------------------ctagtagtgtc-------------

------------------------------------------------------------

------------------------------------------------------------

------------------------------------------------------------

------------------------------------------------------------

------------------------------------------------------------

------------------------------------------------------------

------------------------------------------------------------

---------------------------------------------------------

>Proliferodiscus_sp._TNS__F17436

-------------agcggaggaaaagaaaccaacagggattacctcagtaacggcgagtg

aagcggtaacagctcaaatttgaaatctggctctttcagggtccgagttgtaatttgtag

aagatgctttgggtgtggccccggtctaagttccttggaacaggacgtcatagagggtga

gaatcccgtatgtgactgggtgctttcgcccgtgtaaagctctttcgacgagtcgagttg

tttgggaatgcagctcaaaatgggtggtaaatttcatctaaagctaaatattggccagag

accgatagcgcacaagtagagtgatcgaaagatgaaaagcactttggaaagagagttaaa

cagtacgtgaaattgttgaaagggaagcgcttgcaaccagactcgcatgccgctgatcat

ccggggttctccccggtgcactcggtggtctacgggccagcatcggtttgggtggcggga taaaggctgtgggaatgtagcttctctcggggagtgttatagcccacggtgcaatgccgc

ctacctggaccgaggaccgcgcttcggctaggatgctggcgtaatggttgtaagcgaccc

gtcttgaaa---------------------------------------------------

------------------------------------------------------------

------------------------------------------------------------

------------------------------------------------------------

------------------------------------------------------------

------------------------------------------------------------

------------------------------------------------------------

------------------------------------------------------------

------------------------------------------------------------

------------------------------------------------------------

------------------------------------------------------------

------------------------------------------------------------

------------------------------------------------------------

------------------------------------------------------------

------------------------------------------------------------

------------------------------------------------------------

------------------------------------------------------------

---------------------------------------------------------

>Arachnopeziza_aurelia_TNS__F11211

-------------agcggaggaaaagaaaccaacagggattacctcagtaacggcgagtg

aagcggtaacagctcaaatttgaaatctggctctc---gggcccgagttgtaatttgtag

aagatgctttgggtgtggctccggtctaagttccttggaacaggacgtcatagagggtga

gaatcccgtatgtgactggtcgccttcgcccgtgtaaagctctttcgacgagtcgagttg

tttgggaatgcagctcaaaatgggtggtatatttcatctaaagctaaatattggccagag

accgatagcgcacaagtagagtgatcgaaagatgaaaagcactttggaaagagagttaaa

cagtacgtgaaattgttgaaagggaagcgcttgcaaccagacttgcaggctgttgatcat

ccggggttctccccggtgcactcggcagttttcaggccagcatcggtttgagtggtggga

taaaggccttgggaatgtggctctcttcggggagtgttatagcccaaggtgcaatgccgc

ctacttggaccgaggaccgcgcttcggctaggatgctggcgtaatggttgtaagcgaccc

gtcttg------------------------------------------------------

------------------------------------------------------------
------------------------------------------------------------
------------------------------------------------------------
------------------------------------------------------------
------------------------------------------------------------
------------------------------------------------------------
------------------------------------------------------------
------------------------------------------------------------
------------------------------------------------------------
------------------------------------------------------------
------------------------------------------------------------
------------------------------------------------------------
------------------------------------------------------------
------------------------------------------------------------
------------------------------------------------------------
------------------------------------------------------------
---------------------------------------------------------
>Arachnopeziza_aurata_TNS__F11212
------------agcggaggaaaagaaaccaacagggattacctcagtaacggcgagtg
aagcggtaacagctcaaatttgaaatctggctctc---gggcccgagttgtaatttgtag
aagatgctttgggtgtggctccggtctaagttccttggaacaggacgtcatagagggtga
gaatcccgtacgtgactggtggccttcgctcgtgtaaagctctttcgacgagtcgagttg
tttgggaatgcagctcaaaatgggtggtatatttcatctaaagctaaatattggccagag
accgatagcgcacaagtagagtgatcgaaagatgaaaagcactttggaaagagagttaaa
cagtacgtgaaattgttgaaagggaagcgcttgcaaccagacttgcaggctgttgatcat
ccggggttctccccggtgcactcggcagttttcaggccagcatcggtttgggtggtggga
taaaggccttgggaatgtggctctcttcggggagtgttatagcccaaggtgcaatgccgc
ctacctggaccgaggaccgcgcttcggctaggatgctggcgtaatggttgtaagcgaccc
gtcttg------------------------------------------------------
------------------------------------------------------------
------------------------------------------------------------
------------------------------------------------------------

------------------------------------------------------------
------------------------------------------------------------
------------------------------------------------------------
------------------------------------------------------------
------------------------------------------------------------
------------------------------------------------------------
------------------------------------------------------------
------------------------------------------------------------
------------------------------------------------------------
------------------------------------------------------------
------------------------------------------------------------
------------------------------------------------------------
------------------------------------------------------------
---------------------------------------------------------
>Lachnum_abnorme_KUS__F52080
--------------------------------acagggattacctcagtaacggcgagtg
aagcggtaacagctcaaatttgaaatctggctctttcagggtccgagttgtaatttgtag
aagatgctttgggcgtggctccagtctaagttccttggaacaggacgtcatagagggtga
gaatcccgtatgtgattggtggctttcgcccgtgtaaagctctttcgacgagtcgagttg
tttgggaatgcagctcaaaatgggtggtaaatttcatctaaagctaaatattggccagag
accgatagcgcacaagtagagtgatcgaaagatgaaaagcactttggaaagagagttaaa
cagtacgtgaaattgttgaaagggaagcgttggcaaccagactcgtatgccgctaatcat
ccggggttctccccggtgcacttggtggttttcgggccagcatcggtttcggtggtggga
taaaggccttgggaatgtagcttctctcggggagtgttatagccctcggtgcaatgccgc
ctactgggaccgaggaccgcgcttcggctaggatgctggcgtaatggttgtcaacggccc
gtcttgaaacacggaccaaggagtctaacatctatgcgagtatttgggtgtc-aaaccca
tatgcgtaatgaaagtgaacggaggtaagagccctt-aagggtgcattatcgaccgatcc
tgatgtcttcggatggatttgagtaagagcatagctgttgggacccg-aaagatggtgaa
ctatgcctaaatagggtgaagccagaggaaactctggtggaggctcgcagcggttctgac
gtgcaaatcgatcgtcaaattcgggcataggggcgaaagactaatcgaaccattggaata
cctactaggtggttaagaggcgtaagcctagtcctgcacagggcaacattgtcaaattgt
tcggggacctcccgcacttctcagctaccgcagcctggccgaaaggcgggcgcgcaccag

ggtaacgccctcggggatggtaagaacgctgaaaaggggacgatccgcagcttcttccta

cgggcttcgcctacggagggagttcacagactcgatggcagtgggcctctgggcttaaga

tagagtcgaaccaccgggcaaccggatggagcaatgctagtagctggttcctgccgaag-

---------------tttccctcaggatagcagtgttgtttt------------------

----cagttttatgaggtaaagcgaatgattagaggccttggggttgaaacaaccttaac

ctattctcaaactttaaatatgtaagaagtccttgttacttaattgaacgtggacattcg

aatgtaccaacactagtgggccatttttggtaagcagaactggcgatgcgggatgaaccg

aacgtgaagttaaggtgccggaatct----aggctcatcagacaccacaaaaggtgttag

ttcatctagacagcaggacggtggccatggaagtcggaatccgctaaggagtgtgtaaca

actcacctgccgaatgaactagccctgaaaatggatggcgcttaagcctagtacccatac

ttcaccgccagggtagaaacgatgccctggcgagtaggcaggcgtggaggtcagtga

>Lachnum_palmae_TNS_F__24600

-----------------------------------------gcctcagtaacggcgagtg

aagcggcaacagctcaaatttgaaatctggctccttcggggcccgagttgtaatttgtag

aagatgctttgagggcggc-gcggcctaagttccttggaacaggacgtcagagagggtga

gaatcccgtctggtcaacg----cctaactcgtgtaaagctctttcgacgagtcgagttg

tttgggaatgcagctcaaaatgggtggtaaatttcatctaaagctaaatactggccagag

accgatagcgcacaagtagagtgatcgaaagatgaaaagcactttggaaagagagttaaa

cagtacgtgaaattgttgaaagggaagcgttggcacccagacttgt-cccgaacgctcag

cggggggttcgcccccgtgtattcgtttgtg-gcaggccagcatcggttctggtggcggga

taaa-cccaggggaacgtggctcttc----ggagtgttatagcccttg--gccataccgc

ctaccgggaccgaggaccgcgctttggctaggatgctggcgtaatgggtgtcaacggccc

gtcttgaaacacggaccaaggagtctaacatctatgcaagtatttgggtgctaaaaccca

tatgcgcaatgaaagtgaacggaggtaggcgccctt-aagggtccactatcgaccgatcc

tgatgtcttcggatggatttgagtatgagcatagctgttgggacccg-aaagatggtgaa

ctatgcctaaatagggtgaagccagaggaaactctggtggaggctcgcagcggttctgac

gtgcaaatcgatcgtcaaatttgggcataggggcgaaagactaatcgaactat-------

------------------------------------------------------------

------------------------------------------------------------

------------------------------------------------------------

------------------------------------------------------------

------------------------------------tttgctactaggtagttaagagaa ttataatctagtccctcttgcgtggggcgacactgtcaaattgcggggacgtcctgttat

gctaaactactgcaacattgtggaaacacgtgtgcgcaccagggt---aatgacctgggg

atagtaacaacgtttagagtagggataattcgcagccaagccctaaa----ggttttttg

aacccatgggtgcagttcacagactaaatgtcagtgggcctcgaggcacgctcgaggctt

aagttatagtcggaccgtcggtaaaccgaagagcaagttgggatcttgatagcatgcaag

ctataatattcacgtattatatggctgt-caagagaaatcttgctaaaaactttgtttta

att-------gcgtcgaagcagtccgaaaa------------------------------

------------------------------------------------------caatag

**<u>Alignment of the RPB2 sequences used in the phylogenetic study</u>**

>Lachnum_rachidicola_NBRC_114473

------------------------------------------------------------

------------------------------------------------------------

------------------------------------------------------------

------------------------------------------------------------

------------------------------------------------------------

------------------------------------------------------------

--------tgaaactcccgaaggacaagcttgtgggcttgtcaagaacttggctcttatg

tgttatgtcactgttggtacgcctagtgatcctatcatcgagttcatgatacaaagaaat

atggaagtcctggaagaatatgagcccttgcggtcacctaatgccaccaaagtcttcgtt

aatggcgtttgggttggtgtgcatcgggaccctgcacatcttgtgaggacagtgcagcat

ctgagacgatcacatttgatatctcacgaagtttctttgattcgggatattcgagaccga

gaattcaagatcttcacagacgcaggcagagtgtgcagacctctcttcgtcattgacaac

gacgttgacagtgccaacaagggcaatttggttttgagcaaggagcatatccgacggctt

gaagaagaccaaaccatgcccgcgaatatggactcggaacagaaagccaatgcaggctac

tacggattccaaggcttaattaatgatggtgtggtcgaatatgtggatgctgaagaagaa

gagacagtcatgatcgtgatgacaccggaggacttggatatttctcgtcaactgcaagcg

ggctaccaaatcagaccagatgagagcggagatctgaacaagcgtgtgaaagcgcccatg

aatccaaccgcgcacatatggacccattgcgagattcatccgagtatgattttgggaatt

tgcgcaagcatcattcccttcccggatcacaaccaggtaagacctggagccgatttttgt

tgatcatttcgctgatnattagacagtctcctcgnaacacctacca--------------

-----

>Lachnum_virgineum_TNS__F__16583

------------------------------------------------------------
------------------------------------------------------------
------------------------------------------------------------
------------------------------------------------------------
------------------------------------------------------------
------------------------------------------------------------
-------------------------------gtgggctcgtc-agaacctggctcttatg
tgctacgtcacagtcggtacgcctagcgaccctattattgagttcatgatccaaagaaac
atggaagtcctggaggaatacgagcctttgcggtcgccgaatgcaacgaaagtcttcgtg
aatggtgtatgggttggtgtgcatcgggacccggcacatctcgtgaaaacagtgcagcat
ctgagacgatcacatttaatctcccacgaagtttccctcattcgggatattcgagaccga
gagtttaagatcttcacggacgcaggccgtgtctgtagacctcttttcgtcgttgataac
gatgttgatagtgcgaacaagggcaatttggttttgaacaaggaccacatccgaaaactt
gaagaagatcagacgatgcccgccaatatggacgccgatcagagaactgatgctggatat
tacggtttccagggtttaattaatgatggtgtggtcgagtacgttgatgccgaagaggag
gagactgttatgatcgtaatgactcctgaggacctggatatttcccgtcaattacaagct
ggtttccagatcagaccggatgaaagcggggatctaaacaaacgtgtgaaagcaccgatg
aaccctactgcacatatctggactcattgcgagattcatccaagtatgattttgggaatt
tgcgcaagcattattcccttcccagatcacaatcaggtaaggactagaactattttactc
ggattcctttgctaa---------------------------------------------
-----
>Lachnum_controversum_MFLU_18__1820
--------------------------------------------------tttgttccgc
agattgacgcaggatgtttacaagtacctgacgaaatgtgtttccgagaacaaggaattc
aacctgacacttggagtgaaatttacaaccctcccaaacggtctcaagtactctttggct
actggaaactggggtgatcaaaaaaaagcagctagctcaaccgctggtgtctctcaggtg
ctgaacagatacacgtttgcctccacgctttcgcatttacgccgtacaaatacacctatt
gggcgtgatggaaagatcgccaagccgcgtcaattgcacaatactcactggggtcttgtc
tgtcctgccgaaactcctgaaggacaagcatgtggacttgtcaagaatttggctctcatg
tgttatgtcacagttggtacgcccagcgaccctatcatcgaattcatgatccaaagaaac
atggaagttctagaagaatatgagcctttgcggtctccgaatgcaaccaaagtcttcgtc
aatggtgtgtgggttggtgtgcatcgggaccctgcacacctcgtaaaaacggtgcagcat

ctgcgacgatcacatttgatctctcacgaagtgtctttgattcgtgatattcgagacaga
gaattcaagattttcaccgatgcaggtcgtgtttgcagacctctcttcgttattgacaac
gacgttgacagtactaataagggcaatttggttctgagcaaggagcacatccgacggctt
gaagaggaccagacaatgccggctaatatggactccgaacagaaagctaatgccggctac
tacggtttccaaggcttgattaacgatggtgtagtggaatatgtcgatgccgaagaagag
gagacggtcatgattgtaatgactcctgaggacttggatatctctcgtcaattgcaagct
ggttaccagatcagaccggatgaaagcggggatttgaacaagcgtgtgaaagcaccgatg
aatccaactgcacatatctggacccattgtgagattcatcccagtatgatcttgggaatt
tgcgcaagcattatccccttcccagatcataaccaggtaaggactataattggattccgt
tgaaacttctgc------------------------------------------------
-----
>DSM10201
------------------------------------------------------------
-----------------------gtacctaacgaaatgtgttcaagagaataaggaattc
aacctcacccttggtgtgaaatctacaactctcacaaacggtctcaagtactctctggca
actggaaactggggtgaccagaagaaagcagctagctcgaccgctggtgtgtctcaggtg
ctgaacagatatacatttgcctccacactctcgcatttacgccgtacgaatacgcctatt
gggcgagatggaaagatcgccaaaccgcgtcaactgcacaacactcattggggtctcgtt
tgtcccgccgaaactcccgaaggacaggcttgtgggcttgtcaagaacttggctctcatg
tgttatgtcaccgtgggtactcctagtgaccctattattgagttcatgatccaacgaaac
atggaagttttggaggaatatgagcctttgcggtctccgaatgcgaccaaagtcttcgtc
aatggtgtatgggttggtgtgcatcgggaccctgcacatctcgtaaaaacagtgcagcat
ctgagacgatcacatttgatttctcacgaagtctctttaattcgagatattcgagaccga
gaattcaaaatcttcacagacgcaggccgcgtctgtagacctctcttcgtcattgacaac
gatgttgatagtacaaacaagggaaatttggttttgaacaaggaccacatccgacgactt
gaagatgatcagacgatgcccgcaaatatggactcggaacagaaagccaacgctggttat
tatggtttccagggtttaattaacgatggtgtggtcgagtatgttgatgctgaagaagag
gagactgtaatgatcgtgatgactcctgaggacctggacatttcccgtcaattacaagct
ggttaccagatcagaccggatgaaagtggagatctgaacaagcgtgtgaaagcaccgatg
aatccaactgcacacatctggacccattgcgagatccatccaagtatgattttgggaatt
tgcgcaagcattattcccttcccagatcacaaccaggtaagggctagaattaact-----
------------------------------------------------------------

-----

>ATCC36554

------------------------------------------------------------

------------------------------------------------aataaggaattc

aaccttacccttggtgtgaaatctacaactctcacaaacggtctcaagtactctctggca

actggaaactggggtgaccagaagaaagcagctagctcgaccgctggtgtctctcaggtg

ctgaacaggtatacctttgcctccacactctcgcatttacgccgtacgaatacacctatt

gggcgagatggaaagattgccaagccgcgccaattgcacaacactcattggggtcttgtc

tgtcctgccgaaactcctgaaggacaagcttgtgggcttgtcaagaacttggctctcatg

tgttacgttacagttggtacgcctagtgaccctattattgagttcatgatccaaagaaac

atggaagttctggaggaatatgagcctttgcggtctccgaatgcgaccaaagtcttcgtc

aatggtgtatgggttggtgtacatcgggaccctgcacatctcgtaaaaacagtgcagcat

ctgagacgatcgcatttaatctcccacgaagtttccctgattcgggatattcgagaccga

gagtttaagatcttcacagatgcaggccgtgtctgtagacctcttttcgtcattgacaac

gatgttgatagtgcacagaagggcaatttggttctaaacaaggagcacatccgaaaactt

gaagatgatcagacaatgcccgcaaatatggactcggagcagaaagccaacgctggttat

tatggtttccaaggtttaatcaacgatggtgtggtcgagtacgtcgatgctgaagaagag

gagactgtaatgatcgtgatgactcctgaagacttggatatttcccgtcaattgcaggct

ggctaccagatcagaccggatgaaagtggggacttgaacaagcgtgtgaaagcaccgatg

aatccaactgcacacatctggacccattgcgagattcatccaagtatgattttgggaatt

tgcgcaagcattattcccttcccaga----------------------------------

------------------------------------------------------------

-----

>Lachnum_soppittii_FC__2160

------------------------------------------------------------

------------------------------------------------------------

------------------------------------------------------------

------------------------------------------------------------

------------------------------------------------------------

------------------------------------------------------------

-----------------------------------------------tttggctctcatg

tgttacgtcacagttggtacgcccagcgaccctattattgagttcatgatccaaagaaac atggaagttttggaggaatatgagcctttgcggtctccaaatgcgaccaaagtcttcgtc

aatggtgtatgggttggtgtgcatcgggaccccgcacatctagtgaaaacagtgcagcat

ctgagacgatcgcacttgatctcccacgaagtctccttaattcgggatatccgagaccga

gaattcaagatcttcacagacgcaggtcgtgtctgtagaccactcttcgtcattgacaac

gatgttgagagtaataacaagggcaatttggtcttgaccaaggaccatattcgaagactt

gaagacgatcagacgatgcccgcaaatatggatgcggagcagagagaggcagctaactat

tttggtttccaaggtttaatcaatgagggtgtggtcgagtatgtagatgctgaagaagag

gagactgtcatgatcgtgatgactccagaggacctggacatttcccgccaattgcaagct

gggttgcaaatcagaccagatgaaagcggagacttgaacagacgtgtgaaagcaccgatg

aatccaacagcacacatctggacccattgtgagatccatccaagtatgattttgggaatt

tgcgcaagcattatcccctttccagatcacaaccaggtacggagcagtattgattttggt

tgattcttttgctaatggacatcc------------------------------------

-----

>Lachnum_palmae_TNS_F__24600

------------------------------------------------------------

------------------------------------------------------------

------------------------------------------------------------

------------------------------------------------------------

------------------------------------------------------------

------------------------------------------------------------

tgtcctgctgaaactcctgaaggacaagcttgtggactagtcaagaacctggcactgatg

tgctatgtgacagtcggaacaccgagtgaacccataattgagttcatgatccaaagaaat

atggaagtcttagaagagtacgagcctttacggtctccaaacgcaaccaaagtctttgtt

aatggtgtctgggttggcgtgcaccgcgatcctgctcatcttgtgcgaacagtgcagcat

ctgagacgatcacatctgatctctcatgaagtctctttgatcagagatattcgagataga

gagttcaagattttcaccgatgcaggccgtgtctgtagaccactgttcgtcattgacaac

gatattgacagtgcgaacaaaggtaatttggttttgaataaggagcacatccggcggctt

gaagacgaccaggcgatgcctgccaatatggacgctgatcagagagccagtgctggttac

tatggattccagggcttgatcaatgatggtgttgttgagtacgtggatgctgaggaagaa

gagactgttatgattgtcatgacacctgaagacttggatatttctcgtcagctccaagcc

gggtatcagattagaccagatgagagcggggatctcaataagcgtgtgaaagccccaatg

aacccaactgcacacatctggacgcactgcgagattcatccaagtatgattctagggatt

tgcgcaagcatcatccccttccccgatcacaaccaggtaagaatcaagctaagctttgag
gacgttgctaatgactttagtccc------ctcgtaatacttaccagtctgctatgggta
agcaa
>Lachnum_abnorme_KUS__F52080
------------------------------------------------------------
------------------------------------------------------------
------------------------------------------------------------
------------------------------------------------------------
------------------------------------------------------------
------------------------------------------------------------
------------actcccgagggccaagcttgtggactggtcaagaatttggctcttatg
tgctatgtcacagttggtacgcccagcgacccgattattgagttcatgattcaaaggaac
atggaagtattggaagaatatgaacctctgcggtccccaaatgctacaaaggtctttgtc
aacggcgtttgggttggtgtgcatcgcgatcccgctcatctcgtgaggacagtgcagcat
ctgagacgatcgcatctaatctctcacgaagtttctttgatccgagacattcgtgataga
gaattcaagatctttactgatgcaggccgcgtctgccgaccgctcttcgtcgttgataat
gatgtcgacagtcctaacaaaggcaatttggttttgaataaggagcacatccgacggctt
gaagatgatcaaactatgcctgccaatatggacttagaacagagggctagtgcaggctac
tatggtttccagggtctaatcaatgatggtgtggtcgagtatgtggacgccgaggaagag
gagactgttatgattgtcatgactcccgaggacttggatatttctcgacagttgcaagct
ggttatcagatcagaccagatgaaagcggggatctgaataagcgtgtgaaagcaccgatg
aacccaactgcacatatctggacgcattgcgagattcatccaagtatgatattggggatt
tgcgcaagcattatccccttcccggatcacaaccaggtaataaatccagaacttattcag
agtcccctctctaattgtatc---------------------------------------
-----
>Proliferodiscus_sp._KUS__F52660
aagaagcgtttggatcttgctggacctctccttgctaagctgtttcgaagtctcttccgt
aggctgacaacggatgtctacaagtacctcacgaaatgtgtctctgagaacaaagagttc
aatctcacgctcggagtgaagtccactactctcacgaatggtcttaaatattctttggcc
accgggaactggggtgaccagaagaaagcggcaagctccacggctggtgtctctcaggtg
ctgaacagatatacctttgcttccacactttcgcatttgcgtcgcacaaatacaccaatt
gggcgtgatggaaagattgctaagccgcgacagctgcataacactcactggggtctggtg

tgtcctgctgagactcctgaagggcaagcttgtgggcttgtcaagaatttggctcttatg

tgctacgtcacagtcggtacgcctagcgagcctatcattgagttcatgattcaaagaaat

atggaagttttggaggaatacgagcctctgcggtctcccaatgcaacgaaggtcttcgtc

aacggtgtctgggttggtgtgcaccgcgatcctgcacatctggtgaggacggtgcagcat

ttgaggcgatcgcacttgatctctcacgaagtctctctgattcgagatattcgagacaga

gaattcaagatcttcactgatgcaggccgagtctgtagaccactgttcgtcattgacaat

gatgttgacagtccgaacaaaggcaatctggtcttgaataaggaccacattcgacggctt

gaggatgaccagacgatgcctgccaatatggactcagatcagaaggctaatctaggttat

tatggtttccagggcttgatcaatgatggtgttgttgagtacgttgatgctgaggaagag

gagacagtcatgatcgtgatgactcctgaggacttggacatttctcgccaactgcaagcc

gggtatcagataaggccagatgagagcggcgatctgaacaagcgtgtgaaagcgccaatg

aacccgactgcacacatctggacgcattgcgagattcatcctag----------------

------------------------------------------------------------

------------------------------------------------------------

-----

>Proliferodiscus_sp._TNS__F17436

------------------------------------------------------------

------------------------------------------------------------

------------------------------------------------------------

------------------------------------------------------------

------------------------------------------------------------

------------------------------------------------------------

----------agactcctgaagggcaagcttttgggcttgtcaagaatttggctcttatg

tgctacgtcacagtcggtacgcctagcgagcctatcattgagttcatgattcaaagaaat

atggaagttttggaggaatacgagcctctgcggtctcccaatgcaacgaaggtcttcgtc

aacggtgtctgggttggtgtgcaccgcgatcctgcacatctggtgaggacggtgcagcat

ttgaggcgatcgcacttgatctctcacgaagtctctctgattcgagatattcgagacaga

gaattcaagatcttcactgatgcaggccgagtctgtagaccactgttcgtcattgacaat

gatgttgacagtccgaacaaaggcaatctggtcttgaataaggaccacattcgacggctt

gaggatgaccagacgatgcctgccaatatggactcagatcagaaggctaatctaggttat

tatggtttccagggcttgatcaatgatggtgttgttgagtacgttgatgctgaggaagag

gagacagtcatgatcgtgatgactcctgaggacttggacatttctcgccaactgcaagcc gggtatcagataaggccagatgagagcggcgatctgaacaagcgtgtgaaagcgccaatg

aacccgactgcacacatctggacgcattgcgagattcatcctagtatgattttggggatt

tgcgcaagcattatccccttcccagatcacaatcaggtaagaagatttgtccaatttgcg

tgctcctttactgat---------------------------------------------

-----

>Lachnum_fuscescens_FC__2200

------------------------------------------------------------

------------------------------------------------------------

------------------------------------------------------------

------------------------------------------------------------

------------------------------------------------------------

------------------------------------------------------------

-----------------------------------gcttgtcaagaacctggcgctcatg

tgctacgtcacagtcggtacacccagcgagcctattattgagttcatgatccaaagaaac

atggaggttttggaggagtacgagccgttgaggtccccgaatgccacgaaggtgtttgta

aatggtgtttgggtgggtgtgcacagagaccctgcgcatttggtgaagacagtgcaacat

ctgagacgatcgcatttgatctcccacgaagtctcccttattcgagacatcagagaccga

gaattcaagattttcacagacgcaggacgagtctgcagaccgcttttcgtcattgataac

gatgttgacagtgccaacaagggcaatttggtcttgaacaaggagcacattaggcgactt

gaagaagaccaaactatgccagccaacatggacgccgaacagaaggaaaactccggctac

ttcggattccagggcttgatcgatggtggagtggtcgagtatgtagatgctgaagaagaa

gaaaccgtgatgattgtgatgaccccagaagatttggatatctctcgtcaacttcaggct

ggttaccaaatcagacctgacgatagtggggacttgaacaagcgtgtgaaagcaccgatg

aatcctacagcacatatttggacacattgcgagattcatccaagtatgattttgggaatt

tgcgcaagcattatccctttcccagatcacaatcaagtaaggatctcacagagttt--ga

tgttctttatactaatgtgcc---------------------------------------

-----

>Brunnipila_fuscescens_KUS__F52031

---------ctagatcttgcaggacctcttcttgctaaacttttccgaaatctcttccgc

agattaacaggagatgtgttcaagtatttgcagaaatgtgtggcggataacaaggagttc

aacttgactctgggtgtcaaatccaccaccctcacaaatggtctcaaatattctttggct

accggcaactggggcgaccaaaagaaggcggccagctcaaccgctggtgtgtcccaagtg ctaaacagatacactttcgcttctacactatcccatttacggcgtacaaacacccccatt

ggccgtgatggaaaaattgctaaaccgcgacaacttcacaacactcattggggtcttgtc

tgtcctgccgaaacgcctgaaggacaagcttgtgggcttgtgaagaatttggcgctcatg

tgctacgtcacagtcggtacacccagtgaacctatcattgagttcatgatccaaagaaac

atggaggtcttagaggaatacgagccgttgcggtcgccgaatgccacaaaggttttcgtg

aacggtgtttgggttggtgttcatcgagatccagcgcatctggtgaggacagtgcagcat

ttgagacgatcacatttgatttctcacgaagtttccctgatccgagacattagagatcga

gagttcaagatttttacagatgcaggacgagtctgcagaccgctcttcgtcattgacaat

gatgttgacagtgctaacaagggcaatctggtcttgaacaaggagcacattaggcgactt

gaagaagaccagactatgccagccaacatggaccccgaacaaaaggccaactccggctac

ttcggtttccaaggcttgatcaatgacggtgttgttgaatatgtggacgctgaagaagag

gagactgttatgattgtgatgacccctgaagatttggatatctctcgtcaactacaggcc

gggtaccaaattcgaccagacgatagcggggacctgaacaagcgtgtaaaagcaccaatg

aaccctacagcacacatctggacgcattgcgagatccatcctagtatgattctgggaatt

tgcgcgagcattattcctttcccagatcacaaccaagtaaggagtgaacagagctt--ga

tatatgttttgcta----------------------------------------------

-----

>Trichopeziza_sulphurea_KUS__F52218

------------------------------------------------------------

------------------------------------------------------------

------------------------------------------------------------

------------------------------------------------------------

--------------------------------------gcgaagaacaaacactcctatt

ggtcgagatggaaagattgccaaacctcgtcaattgcacaacactcattggggtctggta

tgtcctgccgagactcctgaggggcaagcttgtggtctggtcaagaatctggcactcatg

tgctatgtcactgttggtacacctagtgacccaatcattgagtttatgatccagagaaac

atggaagttctggaggaatacgaacctttacgttctccaaatgccaccaaggtttttgtg

aatggtgtttgggttggtgtacacagagaccctgcccatcttgtcaggacggtgcaacat

cttcgtcgttctcacttgatctctcacgaagtctccttaattcgagatattcgagatcga

gaattcaaaatcttcacagacgcgggtagagtgtgtagacccttgttcgttattgacaac

gatcttgatagcccaaacaagggaaatctggtactcaataaaatgcacattggacgatta

gaagacgatcaaacaatgcctgcaaatatggacatggaacaaagagtcactcagggtcat

tttggtttccaaggtctcatcaatgaaggtgtggttgaatatgttgatgcggaggaagaa
gagacggtaatgattgtgatgacccccgaagatttggatatctcgcgtcaactccaggca
ggttatcaaattagacctgatgagagtggggatttgaacaaacgtgtgaaagcacctatg
aatccaacagctcatatctggactcattgcgagattcatccaagtatgattctgggaatt
tgcgcaagcattattcctttcccagatcacaatcaggtaagatcattcatgctaagtttc
tg----------------------------------------------------------
-----
>Arachnopeziza_aurelia_TNS__F11211
------------------------------------------------------------
------------------------------------------------------------
------------------------------------------------------------
------------------------------------------------------------
------------------------------------------------------------
------------------------------------------------------------
--------------------------------------------------agctctcatg
tgctacgttacggttggaacgcctagcgatcccattattgagtttatgattcagcgtaac
atggaagttcttgaagaatacgagccactgagatccccaaatgctacaaaagttttcgtc
aacggtgtctgggttggggtccacagagatcctgcccatctggttcagacagtgcaaaat
cttcgaagatctcacttgatctctcacgaggtttcattgattcgagatattcgagaccga
gaattcaagattttcacagatgctggcagagtgtgtcggccacttttcgtcattgaaaat
ggtattgacaatcctaagaaggggcaactagttcttaataaggatcatattcgcagactt
gagcttgatcagacaatg---agtggaatggaccaagatacccgattggctaatggatat
ttcggttttcagggtctaatcaactcgggcgtggttgaatatttggatgccgaagaagaa
gagactgttatgattgtcatgacccctgaggacttagatatctcacgacaactgcaagct
ggatacactcttgaacccgacaccagtggagatatgaataaaagagtgaaggctcctatg
aatcctacagcgcatatgtggacacattgtgagattcatcctagtatgattttgggaatt
tgtgcaagcattattcccttcccagatcataatcaagtaagtacactgt-----------
------------------------------------------------------------
-----
>Arachnopeziza_aurata_TNS__F11212
------------------------------------------------------------
------------------------------------------------------------

------------------------------------------------------------

------------------------------------------------------------

------------------------------------------------------------

------------------------------------------------------------

------------------------------------------------------------

------------------------------------------------------------

------------------------------------------------------------

------------------------------------------------------------

----gaagatctcacttgatatctcacgaagtctctttaatcagagatattcgagaccga

gaatttaaaatcttcacggatgcaggcagagtatgccgacctctttttgttattgaaaat

ggtcatgacaatcccaacaagggaaacttggttttgaacaaagatcatattcgtaagctc

gagttagatcagaccttg---gcaggaatggatcaagaaactcgattggcgaatggatac

tttggcttccaaggtctaatcaactctggagttgtcgaatatctggatgctgaggaagag

gaaacggtcatgatagtcatgactcctgaggatttggatatatcacggcaattacaagct

ggcttcaagattcaacctgacgatagtggggatatgaataagagagttaaggctcctatg

aaccctactgcacatatgtggacgcattgtgagattcatccaagtatgatcttggggatt

tgcgcaagcattatccccttcccagatcataatcaagtaagtccattgttggtcat--ga

tgtctaatgcatgcta--------------------------------------------

-----

Table S1: Reference data for the ten closest matches for the ITS sequence of *Lachnum papyraceum* in GenBank (accessed by a BLAST search on 21 January 2024). The corresponding original literature was retrieved from the Internet if possible.

| Acc no | Taxonomy/Origin | Literature (if available) |
|---|---|---|
| FJ37885 | China, mycorrhizal on *Kobresia* (grass) | Gao,Q. and Yang, Z.L. Ectomycorrhizal fungi associated with two species of *Kobresia* in an alpine meadow in the eastern Himalaya Mycorrhiza 20 (4):281-287 (2010) |
| KF617921 | Unidentified OTU; USA, Alaska, soil under *Picea* | Taylor,D.L., Hollingsworth,T.N., McFarland,J.W., Lennon,N.J., Nusbaum,C. and Ruess,R.W. A first comprehensive census of fungi in soil reveals both hyperdiversity and fine-scale niche partitioning Ecological Monographs (2014) 84(1):3-20 |
| MK808064 | *Lachnum* sp; USA, grassland | Fox,S., Rudgers,J.A., Porras-Alfaro,A et al.. Biogeographic patterns of root-associated fungi in foundation grasses of the North American Great Plains. J. Biogeography 2022 49:22-37 |
| FJ475783 | Helotiales, USA, Oregon, soil | Yarwood SA et al. (2009) Termination of belowground C allocation by trees alters soil fungal and bacterial communities in a boreal forest. FEMS Microbiol Ecol 70:151-162 |
| DQ420921 | Unidentified OTU, USA, soil | Waldrop MP et al. (2006) Resource availability controls fungal diversity across a plant diversity gradient Ecol Lett 9:1127-1135 |
| MK808050 | *Lachnum* sp., USA, grassland | Fox S et al (2022) Biogeographic patterns of root-associated fungi in foundation grasses of the North American Great Plains. J. Biogeography 49:22-37 |
| MT528052 | Unidentified OTU, USA, Alaska, soil | Hewitt RE et al. (2020) Limited overall impacts of ectomycorrhizal inoculation on recruitment of boreal trees into Arctic tundra following wildfire belie species-specific responses. PLoS ONE 15(7): e0235932 |
| MZ158561 | *Lachnum* sp., USA Olean, Cattaraugus County, NY" | Taylor GM (2018) Direct Submission. Specimen depicted on iNaturalist 13413828 |
| MT276007 | *Lachnum pygmaeum,* see discussion in the manuscript text | Ramsfield T et al. (2020) Distance from the forest edge influences soil fungal communities colonizing a reclaimed soil borrow site in boreal mixedwood forest. Forests 11(4):427. |
| OR860210 | *Lachnum sp.,* USA, California, Nov. 2023 | Squazzo,S et al. Direct Submission. Specimen depicted on iNaturalist 165335940 |
